# Evidence that common arbuscular mycorrhizal network alleviates phosphate shortage in interconnected walnut sapling and maize plants

**DOI:** 10.1101/2023.03.08.531711

**Authors:** Emma Mortier, Arnaud Mounier, Jonathan Kreplak, Fabrice Martin-Laurent, Ghislaine Recorbet, Olivier Lamotte

**Affiliations:** Agroécologie, INRAE, Institut Agro Dijon, CNRS, Univ. Bourgogne, Univ. Bourgogne Franche-Comté, 17 rue Sully, BP 86510, F-21065 Dijon, France

**Keywords:** Agroforestry, Phosphate deficiency, Microcosm, Facilitation, *Rhizophagus irregularis*, Symbiotic phosphate transporters, *Juglans* spp.

## Abstract

Under agroforestry practices, inter-specific facilitation between tree rows and cultivated alleys occurs when plants increase the growth of their neighbours especially under nutrient limitation. Due to a coarse roots architecture limiting soil inorganic phosphate (Pi) uptake, walnut trees (*Juglans* spp.) exhibit dependency on soil-borne symbiotic arbuscular mycorrhizal fungi that extend extra-radical hyphae beyond the root Pi depletion zone. To investigate the benefits of mycorrhizal walnuts in alley cropping, we experimentally simulated an agroforestry system in which walnut rootstocks RX1 (*J. regia* x *J. microcarpa*) were connected or not by a common mycelial network (CMN) to maize plants grown under two contrasting Pi levels. Mycorrhizal colonization parameters showed that the inoculum reservoir formed by inoculated walnut donor saplings allowed the mycorrhization of maize recipient roots. Relative to non-mycorrhizal plants and whatever the Pi supply, CMN enabled walnut saplings to access maize Pi fertilization residues according to significant increases in biomass, stem diameter and expression of *JrPHT1;1* and *JrPHT1;2*, two mycorrhiza-inducible phosphate transporter candidates here identified by phylogenic inference of orthologs. In the lowest Pi supply, stem height, leaf Pi concentration and biomass of RX1 were significantly higher than in non-mycorrhizal controls, showing that mycorrhizal connections between walnut and maize roots alleviated Pi deficiency in the mycorrhizal RX1 donor plant. Under Pi limitation, maize recipient plants also benefited from mycorrhization relative to controls, as inferred from larger stem diameter and height, biomass, leaf number, N content and Pi concentration. Mycorrhization-induced Pi uptake generated a higher carbon cost for donor walnut plants than for maize plants by increasing walnut plant photosynthesis to provide the AM fungus with carbon assimilate. Here we show for the first time that CMN alleviates Pi deficiency in co-cultivated walnut and maize plants, and may therefore contribute to limit the use of chemical P fertilizers in agroforestry systems.

## 1. Introduction

Inorganic phosphate (Pi) availability, as the orthophosphate anion PO_4_^-^, is a major factor constraining plant growth and metabolism in many soils worldwide (Bates and Lynch 2001). Consequently, large amounts of Pi fertilizers are used to ensure plant productivity in most conventional agricultural systems (Smith et al. 2011). However, only about 10–30% of the P fertilizer is taken up by the roots, with a substantial part accumulated in the soil as residual P not readily available for plants (Syers et al. 2008). In addition, injudicious and untimely application of chemical fertilizers in agricultural field has generated environmental concerns, including soil degradation and water eutrophication (Tilman 2001; Foley 2005; Conley et al. 2009). To mitigate environmental damages, it is crucial to enhance P-use efficiency in crop production through the diminution of the application of P fertilizers and utilization of residual P and other P pools from soils (Xue et al. 2016). As a substitute to conventional cropping systems, which rely on large inputs of chemical fertilizers to sustain production, agroforestry is a low-input design which combines trees with annual crops in various combinations or sequences (Nair 1993). Agroforestry systems aim at reducing the need for inputs through minimising losses and maximising internal cycling of nutrients (Smith 2010). One of the major arguments in favour of agroforestry is the fact that a mixed system of trees and annual crops make a better use of natural resources such as light, water and nutrient more efficient than a monoculture. As resources are used more efficiently, less chemical fertilizers are required to the benefit of the overall environment (Postma 2005).

A fundamental aspect of agroforestry is to favour inter-specific facilitation between tree rows and cultivated alleys, where plants increase the growth and survival of their neighbours (Callaway 1998), particularly regarding limited resources such as nutrients (Schoeneberger et al. 2012; Battie-Laclau et al. 2020). In the maintenance of soil P fertility under alley cropping systems, the role of roots is at least as important as that of aboveground biomass (Young 1997; van Noordwijk et al. 2015). Belowground facilitation occurs when one species makes previously unavailable P available to the other, due to the exudation of organic acids, phosphatases, or rhizosphere acidification resulting in increased P availability (Hinsinger et al. 2011). Since a decade ago, numerous studies have also pointed to the importance of soil-borne arbuscular mycorrhizal (AM) fungi of the phylum Glomeromycota (Tedersoo et al. 2020) in mediating inter-plant facilitation processes (van der Heijden et al. 2008; Montesinos-Navarro et al. 2016). In soils with strong Pi-fixing capacity, or where Pi is limiting, plant demand for this nutrient exceeds the rate at which it diffuses into the root zone, resulting in zones of Pi depletion surrounding the roots (Smith and Read 1997). AM fungi help to overcome this limitation by extending their extra-radical hyphae from root surfaces to areas beyond the P depletion zone (Hayman 1983; Javot et al. 2007). When roots are colonized by AM fungi, soil Pi is acquired at the soil-fungus interface through high-affinity fungal phosphate transporters (PHTs) located in the extra-radical mycelium (Rui et al. 2022). In the plant interfacial apoplast of fungal arbuscules, Pi is acquired by plant PHT located on the peri-arbuscular membrane (Harrison et al. 2002; Javot et al. 2007). The Pi uptake pathway of mycorrhizae may dominate Pi uptake in AM symbiosis, which is heavily dependent on AM-induced PHT1 members (Casieri et al. 2013; Rui et al. 2022). Consequently, AM fungi have the potential to maintain crop yields at low soil P levels through a more effective exploitation of available P sources. It has been estimated that inoculation with AM fungi might reach a reduction of approximately 80% of the recommended fertilizer P rates under certain conditions (Jakobsen et al. 1992). In return, AM fungi are supplied with sugar and lipid from the host plant (Gutjahr and Parniske 2013; Roth and Paszkowski 2017). Conversely, the importance of the role of mycorrhizal symbiosis in providing Pi to the plant usually declines when soil phosphate concentration is elevated (Kiers et al. 2011).

As many other tree species used in tropical and temperate alley cropping systems such as ash, cocoa, coffee, eucalyptus, olive tree, paulownia, poplar, and rubber (Cardoso 2002; Janos 2007; de Kroon et al. 2012), walnut (the common name given to species of deciduous trees belonging to the genus *Juglans*) exhibits a high dependency on symbiotic AM fungi for their development. Namely, coarse roots and relatively few roots per plant limit soil Pi uptake (Brundrett 1991; Comas and Eissenstat 2009; Comas, et al. 2014). By providing wood, ornamental, and nutrition value to human beings, and food and a habitat to wildlife, walnuts belong to the most important trees in the northern hemisphere, ecologically and economically (Bernard et al. 2018). Because of its short growing season, sparse canopy and deep rooting system, walnut is an ideal species for agroforestry and belongs to the dominant trees used in temperate alley cropping (Wolz and DeLucia 2018). Intercropping of *Juglans* spp. is aimed at increasing growth and quality of walnut trees together with providing an early financial return to help offsets the costs associated with establishing walnut orchards (van Sambeek and Garrett 2004; Mohni et al. 2009; Wolz and DeLucia 2019). Before walnut plantations reaches economic maturity, the intercrop - winter cereals (*Triticum* spp), alfafa (*Medicago sativa*), soybean (*Glycine max*), or summer crops (e.g., maize) - is the only income during the first five to ten years; thereafter both trees and intercrops produce simultaneously (Mary 1998). Among these AM-dependent intercrops, maize (*Zea mays* L.), is one of the most cultivated crops for both staple food and industrial usage, in tropical and temperate soils worldwide, with phosphorus being a major growth limiting factor for commercial maize production (Brady and Weil 2008).

The role of AM fungi in mixed plant communities is applicable due to their low host specificity: extra-radical hyphae can connect the roots of different plant species to form a common mycelial network (CMN) (Ingleby et al. 2007). This inter-plant mycorrhizal connection promotes facilitation through differential access to the nutrient pool from the common hyphal network and plant-to-plant nutrient transfers (Martins and Cruz 1998; Fitter et al. 1998; Leake et al. 2004; Hauggaard-Nielsen and Jensen 2005; van der Heijden and Horton 2009; Walder et al. 2012; Fellbaum et al. 2014; Gorzelak et al. 2015; Montesinos-Navarro et al. 2016). Remarkably, when associated to deep perennial tree roots, AM fungi increase the mycorrhiza-mediated uptake of soil nutrients, and thereby contribute to the nutrition of the co-cultivated species (Simard and Durall 2004; Ingleby et al. 2007; de Carvalho et al. 2010; Bainard, et al. 2011; Bainard et al. 2012). In parallel, when focussing on the effect of intercrops on tree growth, tree display a larger growth than that observed in simple forestry systems, due to a better mineral nutrition even in low input systems (Jose et al. 2000). To date, improved tree growth has been ascribed to the likely ability of trees roots to intercept and uptake a significant proportion of nutrient fertilizer residues from the crop rooting zone (Smith 2010), but the contribution of inter-plant mycorrhizal connections to this process yet remains poorly investigated. Because improving tree mineral nutrition through agroforestry systems is a low-cost means to increase tree growth and to shorten the development duration before tree harvesting (Chifflot 2009), this information is of importance to assess the overall costs or benefits of growing trees and annuals.

In this study, we investigated for the first time the extent to which intercropping with maize contributes to the mineral nutrition and biomass production of young hybrid walnut plants through the formation of a common AM fungal mycelial network. To this aim, we experimentally simulated an agroforestry system using a home-made experimental device in which young walnut hybrid rootstocks were connected or not by a common mycelial network to maize plants grown under contrasting Pi supply levels (0.13 *vs* 1.3 mM). This experimental device was made of two separated compartments in which walnut and maize were planted and connected by a third compartment in which the extra-radical mycelium responsible for mineral nutrient supply for the plants, was separated by fine nylon nets from the associated host roots (**Fig. 1**). Two months after plant sowing, we analysed plant growth parameters, nutrient contents, and expression of plant genes coding mycorrhiza-inducible PHT1 transporters, here identified by phylogenic inference of orthologs. We hypothesized that (1) the inoculum reservoir formed by walnut roots would allow the mycorrhization of maize roots through the formation of a CMN, thereby giving access to maize fertilization residues; (2) hyphal connections under P deficient condition would result in a benefit comparable to P sufficient condition without a CMN; and (3) due to a slower growth rate than maize, walnut would invest more carbon in the development of the mycorrhizal fungus than maize.

**Figure 1.**
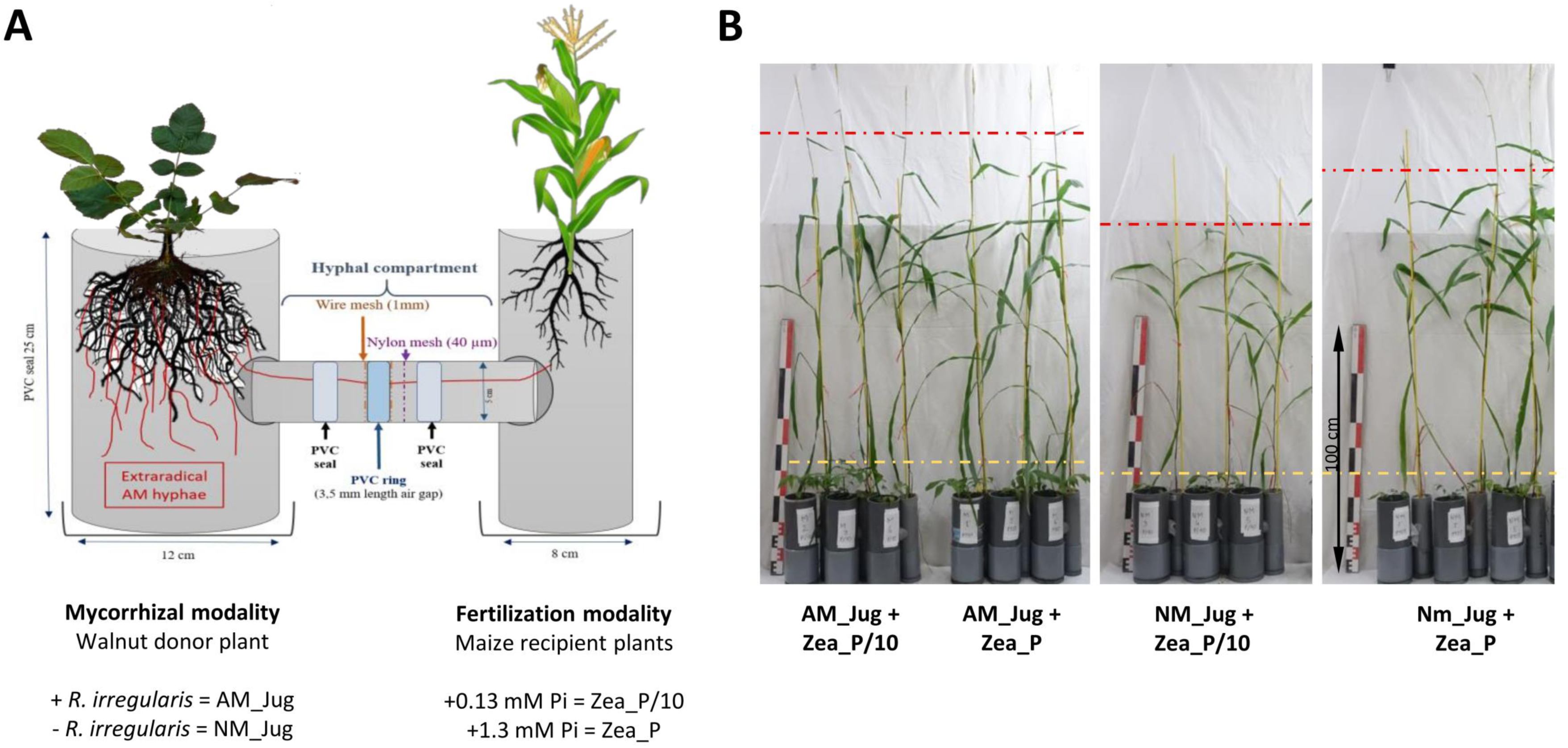
Schematic representation (**A**) and pictures (**B**) of the experimental device in which young walnut hybrid roostocks (Jug) were connected (AM) or not (NM) by a common mycelial network to maize plants (Zea) grown under two contrasting Pi supply levels (Zea_P/10 and Zea_P). Yellow and red dashed lines show the stem height reached by walnut and maize plants, respectively. Pictures in panel B were taken two months after plant sowing in microcosms.

## 2. Materials and Methods

### 2.1 Biological material

Micropropagated plantlets of the walnut hydrid rootstock RX1 (*J. regia x J. microcarpa*) (Browne et al. 2015) were provided by Dr. Laurent Jouve (Tarnagas, Brive-la-Gaillarde, France). Rootstocks were received seven weeks after root induction and placed for *ex vitro* acclimatization in closed mini-greenhouses (Rapid Grow, Nortene) filled with a sterilized (120 °C, 12h) quartz sand (Biot, France): zeolite (Symbion, Czech Republic) substrate (1:1, v:v). They were sprayed twice a week with tap water to maintain humidity and fertilized once a week with 15 ml per plant of a NPK 6- 4-6 solution containing 2% of magnesium and oligoelements (Algoflash, France). Walnut cultures were maintained in a growth chamber under controlled conditions (16 h photoperiod, 23 °C/18 °C day/night, 100% relative humidity, 220 µE m-2 s-1). After two months of growth, plantlets were gradually exposed to reduced relative humidity by progressively removing the box covers during a further month, then RX1 rootstocks were planted in microcosms.

Maize (*Zea mays* L. var. Pompeo) seeds were surface-disinfected for 15 min in a commercial solution of sodium hypochlorite (1% available chlorine), and rinsed thoroughly. After imbibition in sterile water for 12 h in the dark at 4 °C, seeds were germinated at 20 °C with 16 h of fluorescent light (Osram Fluora L36W/77) on filter papers moist with sterile water. Maize plantlets were sown in microcosms at the third leaf stage.

The isolate DAOM197198 (also known as DAOM181602) of the model AM fungus *Rhizophagus irregularis* originally isolated from Pont Rouge, Canada (Stockinger et al. 2009) and commercialized by Agronutrition (Labège, France) was purchased as an axenic spore suspension (200 spores/ml).

### 2.2 Microcosm set-up, plant inoculation, and phosphate fertilization

Microcosms were set-up in a home-made device comprising tripartite compartments consisting of two root hyphal compartments, each connected to one central hyphal pipe (**Fig.1**). Root hyphal compartments were made out of PVC tubes, each having a hole cut to allow the insertion of the horizontal central PVC tube (7.5 cm length, 5 cm inner diameter) containing the hyphal compartment. The walnut mycorrhizal donor plant compartment was designed wider (25 cm height, 12 cm inner diameter) than the maize plant receiver compartment (25 cm height, 8 cm inner diameter) to allow maize plants to explore beyond their own compartment (Teste et al. 2015). To create an air gap, a PVC ring (3.5 cm length, 5 cm external diameter) was vertically inserted in the central PVC tube and distributed at equal distance from each root hyphal compartment. In order to prevent a substrate-mediated diffusion of solutes between both sides of the PVC ring, the air gap was sandwiched between two 1-mm wire meshes using two PVC seals. Both sides of the air gap had a 40-µm nylon membrane (Anachem, Bedfordshire, UK) through which hyphae, but not plant roots, can grow (Wang et al. 2016). Two additional plastic boxes were placed to the base of each of the root hyphal compartments to collect drainage solutions. Root hyphal and central compartments were filled with the corresponding volume of a sterilized (120 °C, 12h) quartz sand (Biot, France): zeolite (Symbion, Czech Republic) substrate (1:1, v:v). Walnut rootstocks were planted into the wider root hyphal compartment and maize seedlings into the other one. To assess whether mycorrhizal walnuts can benefit maize plant growth through a CMN, four treatments were compared. For the mycorrhizal modality, to check whether inoculated saplings in the nursery should serve as infection sources for the interrow crop via a CMN, only donor walnut plants were inoculated (AM_Jug) or not (NM_Jug) with 1 ml of the spore suspension of *R. irregularis*. For the Pi modality, only recipient maize plants were fertilized in order to simulate nutrient fertilizer residues in the crop rooting zone. To this aim, maize plant received once a week 10 ml of either a low-Pi (Zea_P/10: 0.13 mM NaH_2_PO4, 2H_2_O) or a full strength Pi solution (Zea_P: 1.3 mM NaH_2_PO4, 2H_2_O) Hoagland solution (Bonneau et al. 2013), while walnut donor saplings received 10 ml tap water. Plants were maintained in a growth chamber under controlled conditions with 16 h of light [220 µE m**^-^**^2^ s**^-^**^1^] at 23 °C and 8 h of dark at 19 °C for two months before harvest. For each treatment, the experimental setup was replicated at least six times.

### 2.3 Measurement of plant growth and nutritional parameters

Before harvest, the leaf chlorophyll *a* fluorescence was quantified on attached donor and recipient leaves using a Pulse-Amplitude-Modulation (PAM) Chlorophyll Fluorimeter (Junior PAM, Walz, Effeltrich, Germany). Plants were dark-adapted for at least one hour and measurements were further performed on leaf adaxial side. From these measurements, the maximum photochemical quantum yield of PS_II_ (Fv/Fm) and the effective photochemical quantum yield of PS_II_ (Y_II_) were calculated as described in the operator’s guide. For plant growth parameters, plant height (cm), branch or leaf number per plant, stem diameter at the soil line (mm), fresh shoot and root weights (g), and primary root length (cm) were measured after two months after sawing. To determine dry matter weights (DW), shoot and root subsamples were separately weighed fresh, and subsequently dried for 2 days at 80 °C and weighed. For soluble Pi, C and N measurements, dried root and leaf samples were finely ground using a Retsch Mixer ball mill (Haan, Germany). Soluble Pi was extracted from 5 mg dried samples as described by Grunwald et al. (2009). Free Pi was determined by measuring absorbance at 650 nm using the BIOMOL Green™ Reagent (Enzo Life Sciences, Villeurbanne, France) according to the instructions of the manufacturer. Root and leaf dried aliquots (5 mg) were analysed for their C and N contents at the Biogeosciences Laboratory of the University of Bourgogne Franche-Comté (Dijon, France) on a FlashSmart Elemental Analyzer (Thermo Scientific).

### 2.4 Staining of AM fungal propagules and quantification of root colonization

At harvest, root samples were randomly collected from each plant and dried for 2 days at 80 °C (Hetrick et al. 1989). After overnight rehydratation in water (Tarkalson et al. 1998), they were cleared with at 90 °C for 30 min in 10% (w/v) KOH (Phillips and Hayman 1970). Walnut roots were additionally bleached with three drops of 30% H_2_O_2_ to remove any phenolic compounds left in cleared roots (Kormanik and McGraw 1982). Roots were rinsed with tap water before staining in a 0.05% (w/v) trypan blue (Sigma-Aldrich) solution in lactoglycerol (lactic acid-glycerol demineralised water, 1:1:1) for 30 min at 90 °C. Thirty fragments, ca. 10 mm long, were randomly collected from each sample, mounted 15 per slide in glycerol and observed at a magnification of 200 as described in Trouvelot et al. (1986). Calculations of frequency of mycorrhization (F%), percentage of root cortex colonisation (M%) and percentage of arbuscules (A%) were carried out with the MycoCalc program (http://www.dijon.inra.fr/mychintec/Mycocalc-prg/download.html). Similarly to roots but without drying, clearing, and bleaching steps, the 40-µm nylon mesh and the substrate collected in the half-hyphal pipes connecting the maize compartment were also stained in trypan blue for 30 min at room temperature and rinsed with tap water before examination.

### 2.5 Identification and *in silico* characterization of proteins orthologous to the AM-inducible phosphate transporter MtPT4 of *Medicago truncatula* in *J. regia*, *J. microcarpa* and *Z. mays*

To identify walnut RX1 and maize plant PHT1 transporters orthologous to the AM-inducible phosphate transporter MtPT4 of the model legume *Medicago truncatula* (Harrison et al. 2002), we used Orthofinder (version 2.2.6; Emms and Kelly 2015) through standard mode parameters in the Diamond tool (v0.9.22.123) all-versus-all. To this aim, the complete predicted proteomes of *J. regia*, *J. microcarpa*, *M. truncatula,* and *Z. mays* were download at: https://treegenesdb.org/FTP/Genomes/Jure/v2.0/annotation/Jure.2_0.pep.fa.gz, https://treegenesdb.org/FTP/Genomes/Jumi/v1.0/annotation/jumi.1_0.peptides.fa, https://genome.jgi.doe.gov/portal/pages/dynamicOrganismDownload.jsf?organism=Mtruncatula/Mtruncatula_198_protein.fa.gz and https://ftp.ensemblgenomes.org/pub/release-43/plants/fasta/zea_mays/pep/Zea_mays.B73_RefGen_v4.pep.all.fa.gz, respectively. Bioinformatics tools available at NGPhylogeny.fr (https://ngphylogeny.fr) were used according to the one click mode with MAFFT for multiple alignment, BGME for alignment refinement, FastME for tree inference, Newick Display for tree rendering, and iTOL for phylogenic tree display and annotation (Lemoine et al. 2019).

Protein sequences were aligned using MultAlin (http://multalin.toulouse.inra.fr/multalin/) Alpha-helical trans-membrane (TM) regions together with cyto-and exo-plasmic sequence orientations were predicted according to DeepTMHMM (https://dtu.biolib.com/DeepTMHMM).

### 2.6 Primer design

On the basis of the plant orthologous proteins identified using Orthofinder, corresponding nucleotide sequences were retrieved to design oligonucleotide primers with Primer-BLAST (https://www.ncbi.nlm.nih.gov/tools/primer-blast). To monitor Pi acquisition through the fungal side, expression pattern of the gene coding the high affinity transporter RiPT1 previously found to be regulated by external Pi concentrations in *R. irregularis* (Calabrese et al. 2019) was monitored in mRNA extracts of walnut and maize roots. All primers used are listed in **Supplementary Table 1**.

### 2.7 mRNA extraction, reverse transcription and qRT-PCR

At harvest, maize and walnut root subsamples (1.5 g) were immediately frozen in liquid nitrogen and kept at −80 °C before use. They were ground to a fine powder with a pre-cooled mortar and pestle before mRNA extraction. The SV Total RNA Isolation System (Promega, Madison, USA), which incorporates a DNase treatment step, was used to recover total root RNA from 50 mg ground tissues, according to the supplier instructions. For walnut, 50 mg of insoluble polyvinylpyrrolidone were added to root samples before mRNA extraction in order to remove polyphenols (Hu et al. 2002). Purified mRNAs were quantified by measuring the absorbance at 260 nm with a Nanodrop (ND-1000 spectrophotometer, Isogen Life Sciences, Maarssen, Netherlands). Complementary DNAs (cDNAs) were obtained using High Capacity cDNA Reverse Transcription Kit (Thermo Fisher Scientific) using 300 ng of total mRNA per reaction. Quantitative RT-PCRs were run in a 7500 real-time PCR system (Roche) using the following settings: 95°C for 3 min and then 40 cycles of 95 °C for 30 s, 60 °C for 1 min and 72 °C for 30 s. These analyses comprised at least five biologicals and three technical replicates per treatment. Transcript levels were expressed as normalized expression to previously described reference genes (**Table S1**).

### 2.8 Statistical analyses

For each plant, mycorrhizal colonization parameters were compared between the two fertilization regimes with the Welch’s t-test (unequal variances t-test), after *arcsin square root* transformation of percentages. For comparison of growth, nutritional and gene expression parameters between treatments, the Kruskal-Wallis H-test with post-hoc Tukey HSD (Honestly Significant Difference) were used under the Rstudio environment (version 4.03) due to non-Gaussian distribution of data and/or inequality of variances, as checked by the the Kolmogrov–Smirnov’s and and Levene’s tests, respectively. *P*-values less than 0.05 were used for statistical significance.

## 3. Results

### 3.1 Mycorrhizal colonization and extra-radical hypha development

In walnut roots inoculated with *R. irregularis*, the frequency of colonization (F%), the intensity of root cortex colonization (M%) and arbuscule abundance (A%) in the Zea_P/10 treatment reached 100, 20 and 10%, respectively (**Fig. 2A**). The intensity of root cortex colonization and arbuscule abundance in AM-Jug roots were significantly (*p* < 0.05) higher in the Zea_P/10 treatment when compared to Zea_P (**Fig. 2A**). In non-inoculated maize roots connected to AM walnut plants, F, M, and A% values reached 100, 30 and 25%, respectively, in both Pi fertilization regimes (**Fig. 2B).** Whatever the fertilization regime applied to maize plants, extra-radical hyphae were visualized after trypan blue staining of the nylon mesh (**Fig. 2C**) and the substrate (**Fig. 2D**) contained in the hyphal pipes close to the maize root compartment. In both Pi fertilization regimes, walnut and maize plants were not colonized when the walnut donor plant was not inoculated (negative control).

**Figure 2.**
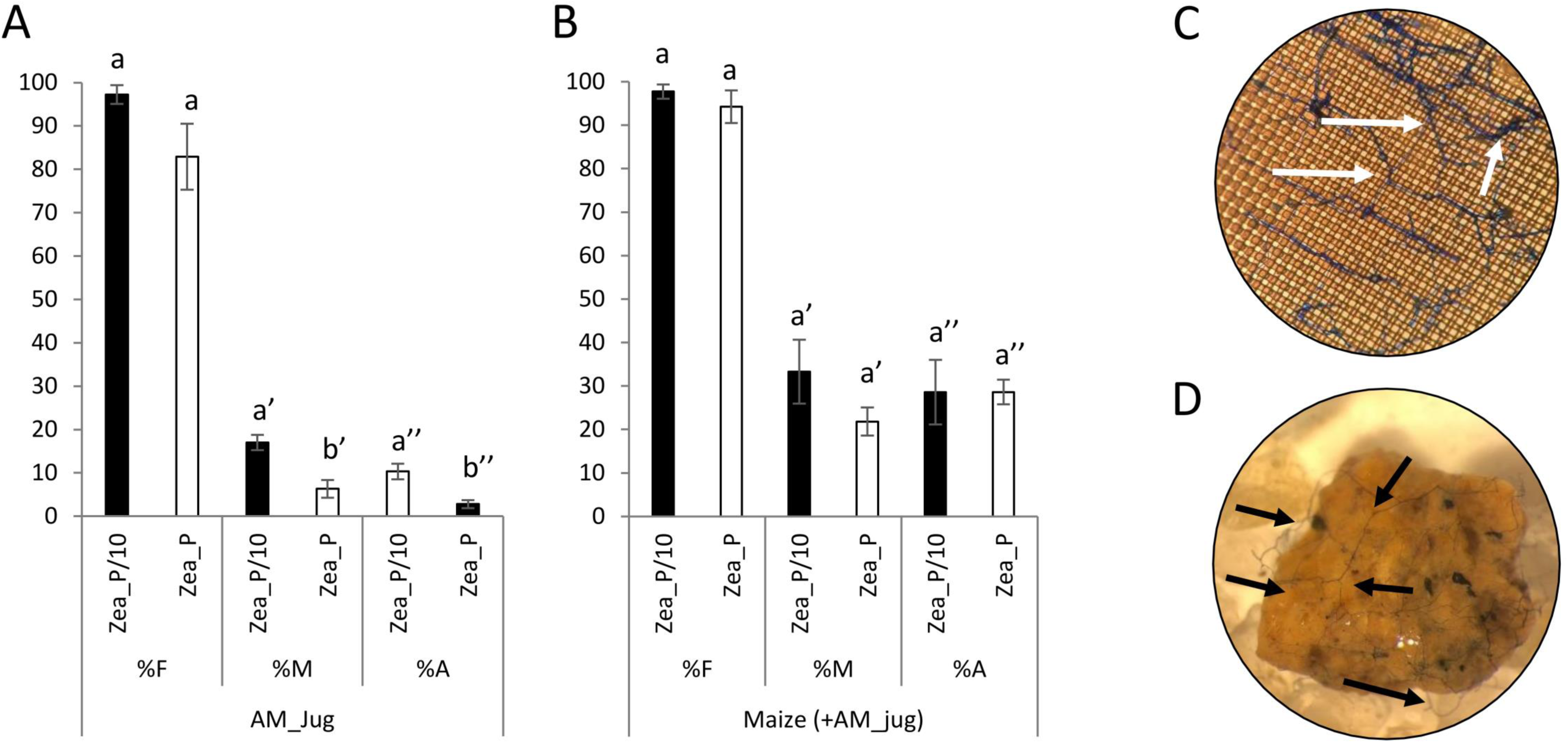
Mycorrhizal colonization parameters measured two months after inoculation of walnut RX1 plants with *R. irregularis* under contrasting Pi fertilization regimes of maize (Zea_P/10, Zea_P) in (**A**) walnut donor and, (**B**) non-inoculated maize recipient roots. Values correspond to the mean (±SE) of six replicates per treatment. Lower case letters indicate significant difference (Welch’s t-test after arcsin square root transformation; *p* < 0.05). Whatever the Pi fertilization regime of maize, white and black arrows show the development of the extra-radical mycelium on the nylon mesh (**C**) and the substrate (**D**) in the hyphal compartment close to maize roots after trypan blue staining and observation at a magnification of 100 and 16, respectively.

### 3.2 Effect of walnut mycorrhization and maize Pi fertilization on rootstock growth and nutritional parameters

In walnut roots, in both Pi-limited (P/10) and Pi-replete (P) fertilization regimes of maize, mycorrhization of walnut (AM_Jug) led to a significant increase (*p* < 0.05) in root collar, root and shoot dry weights (DW), relative to non-mycorrhizal (NM_Jug) plants (**Fig. 3A**). Under the maize Pi-limited condition, but not in the Pi-replete condition, AM walnut plants displayed a higher (*p* < 0.05) stem height, leaf Pi concentration and effective photochemical quantum yield of PSII (Y_II_), together with a decreased root carbon content and shoot to root biomass ratio when compared to NM saplings (**Fig. 3B**). All the other parameters measured in walnut plants (the maximum photochemical quantum yield of PS II (Fv/Fm), primary root length, the number of branches, root soluble Pi, leaf C content, plant N content, and leaf C/N ratio) were not significantly modified either by walnut mycorrhization or Pi fertilization regime applied to maize plants (**Supplementary Table 2**).

**Figure 3.**
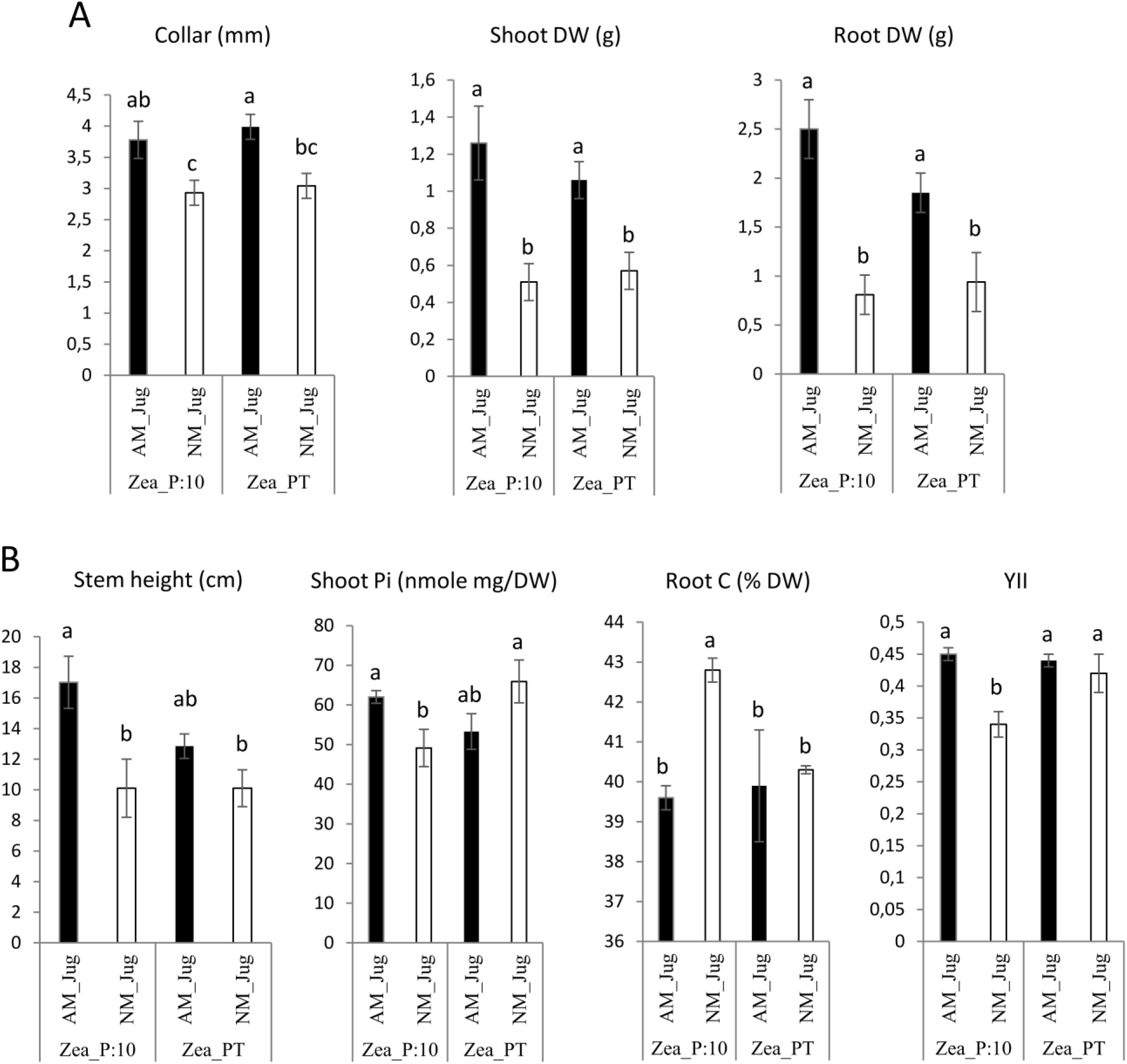
Effect walnut mycorrhization (AM) as compared to non-mycorhizal controls (NM), on the growth and nutritional parameters of RX1, irrespective of Pi fertilization of *Z. maïs* (**A)**, or only recorded in the P/10 condition (**B**). Values correspond to the mean (±SE) of six replicates per treatment. Lower case letters indicate significant difference (Kruskal-Wallis H-test with post-hoc Tukey HSD; *p* < 0.05).

### 3.4 Effect of walnut mycorrhization and maize Pi fertilization on *Z. mays* growth and nutritional parameters

In maize plants, walnut mycorrhization with *R. irregularis* (AM_Jug) as compared to NM plants, led to a significant increase in root collar diameter, height, leaf number, root and shoot DW, leaf N content and Pi concentration, and to a decrease in leaf C/N ratio under the maize Pi-limited condition, but not in the Pi-replete condition (**Fig. 1**, **Fig. 4**). The other parameters measured in maize plants namely Y_II_, Fv/Fm, primary root length, shoot to root biomass ratio, root Pi, C and N root and leaf contents, were not significantly modified by the mycorrhization of walnut with *R. irregularis* (**Table S2**). As regards Pi availability, the maximum photochemical quantum yield of PS_II_ (Fv/Fm) was significantly but slightly decreased only in NM plants under the low-Pi condition, when compared to the Pi-replete fertilization regime (**Table S2)**.

**Figure 4.**
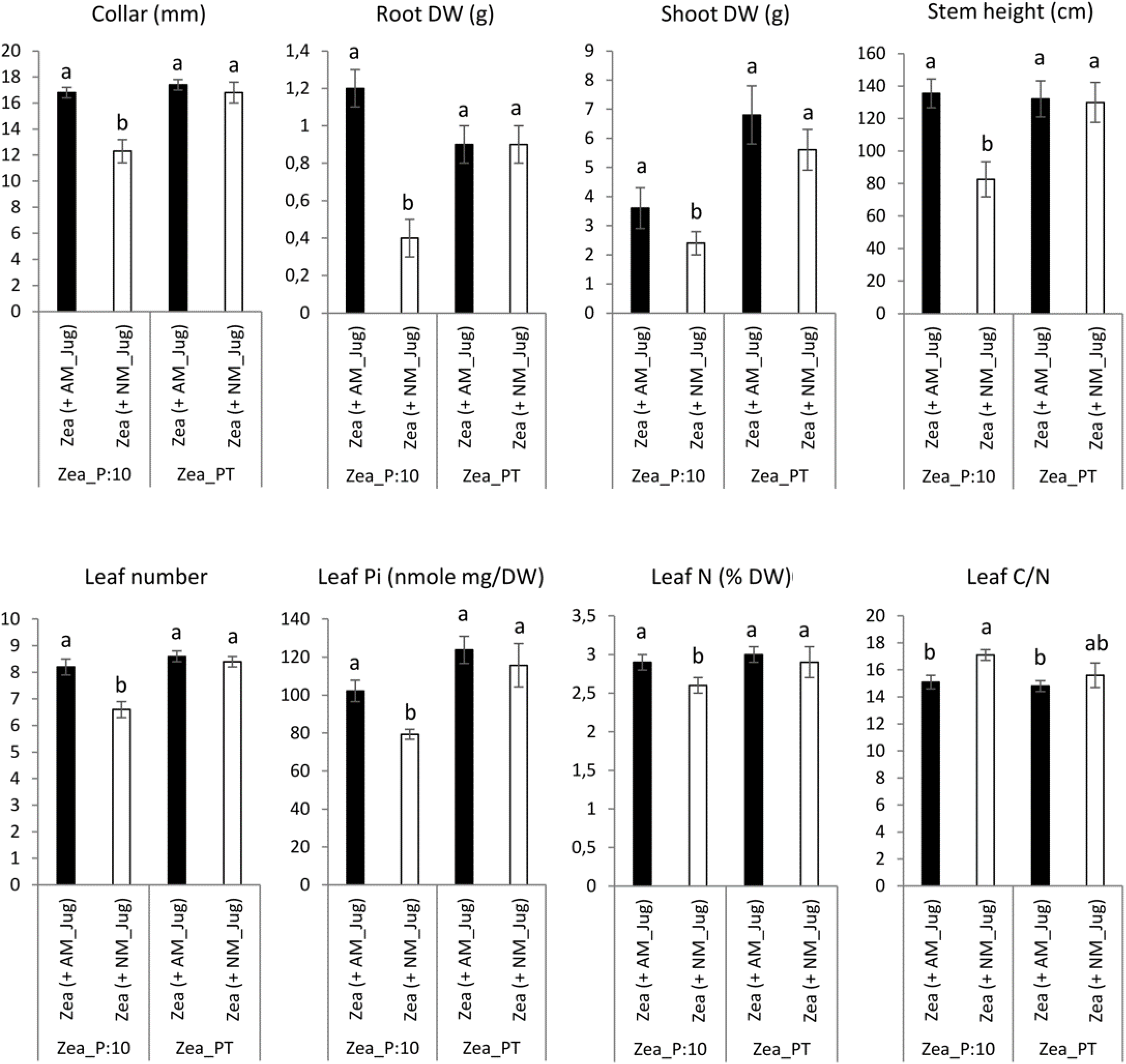
Effect of walnut mycorrhization (AM) as compared to non-mycorhizal controls (NM), on the growth and nutritional parameters of maize plants only recorded under the P/10 condition. Values correspond to the mean (±SE) of six replicates per treatment. Lower case letters indicate significant difference (Kruskal-Wallis H-test with post-hoc Tukey HSD; *p* < 0.05).

### 3.5 Identification of walnut and maize proteins orthologous to the AM-inducible phosphate transporter MtPT4 of *M. truncatula*

To identify walnut RX1 (*J. regia* x J*. microcarpa*) and maize plant PHT1 transporters orthologous to the AM-inducible phosphate transporter MtPT4 of *M. truncatula*, we performed an OrthoFinder analysis that included the predicted proteomes of *J. microcarpa*, *J. regia*, *Z. mays*, and the reference proteome of *M. truncatula*. OrthoFinder outputs (**Supplementary Table 3; Supplementary Fig. 1**) indicated that the MtPT4 orthogroup contained three proteins of *J. microcarpa* (Jumi_00855.t1, Jumi_15132.t1, Jumi_21692.t1, further named JmPHT1;1, JmPHT1;2, JmPHT1;3, respectively), three proteins of *J. regia* (Jr13_30200_p1, Jr13_30210_p1, Jr16_00830_p1, further named JrPHT1;1; JrPHT1;2, JrPHT1;3, respectively), and one protein of *Z. mays* (Zm00001d011498_P001). The latter protein turned out to be the AM-inducible Pi transporter ZmPHT1;6 previously described in maize (Glassop et al. 2005; Tian et al. 2013). **Fig. S1** shows that all the seven proteins retrieved in the MtPT4 orthogroup display the PHT1 signature consisting of the conserved sequence GGDYPLSATIxSE of 12 amino acid residues (Karandashov and Bucher 2005). Hydrophobicity analysis (**Fig. S1**) further indicated that they also share a common topology with PHT1 transporters, as inferred from 12 membrane-spanning domains, which are separated into two groups of six domains by one intracellular central loop to transfer the phosphate molecule (Poirier and Bucher 2002). As expected, both C- and N-termini were predicted to be oriented inside the cell (Nussaume et al. 2011). According to the nomenclature of Naguy et al. (2005), the phylogenetic tree of the PHT1 protein family built with 81 additional amino acids sequences with substantial diversity and redundancy within dicots and monocots (**Table S4**), showed that all the protein candidates retrieved as MtPT4 ortologs in *J. regia* and *J. microcarpa*, are associated to the AM-inducible subfamily I (**Fig. 5**). The latter cluster includes the mycorrhiza-specific Pi transporters OsPT11, which similarly to MtPT4 (Pumplin and Harrison 2009), has been localized at the arbuscule-branch domain of the peri-arbuscular membrane (Kobae and Hata 2010). Reduced fungal growth and altered arbuscule morphology in *Medicago pt4* and rice *pt11* loss-of-function mutants also indicated that PT4 and PT11 are essential for AM-mediated phosphate uptake and arbuscule maintenance (Javot et al. 2007; Yang et al. 2012). Likewise, ZmPt1;6 is a mycorrhiza-specific Pi transporter of maize, and a knock-down *pht1;6* mutant showed reduced mycorrhiza formation (Willmann et al. 2013). In dicot plant species, the other AM-inducible Pi transporters that cluster in subfamily III (**Fig. 5**) such as *StPT3* in potato (Rausch et al. 2001), are over-expressed in mycorrhizal roots but are also expressed in non-symbiotic conditions (Casieri et al. 2013).

**Figure 5.**
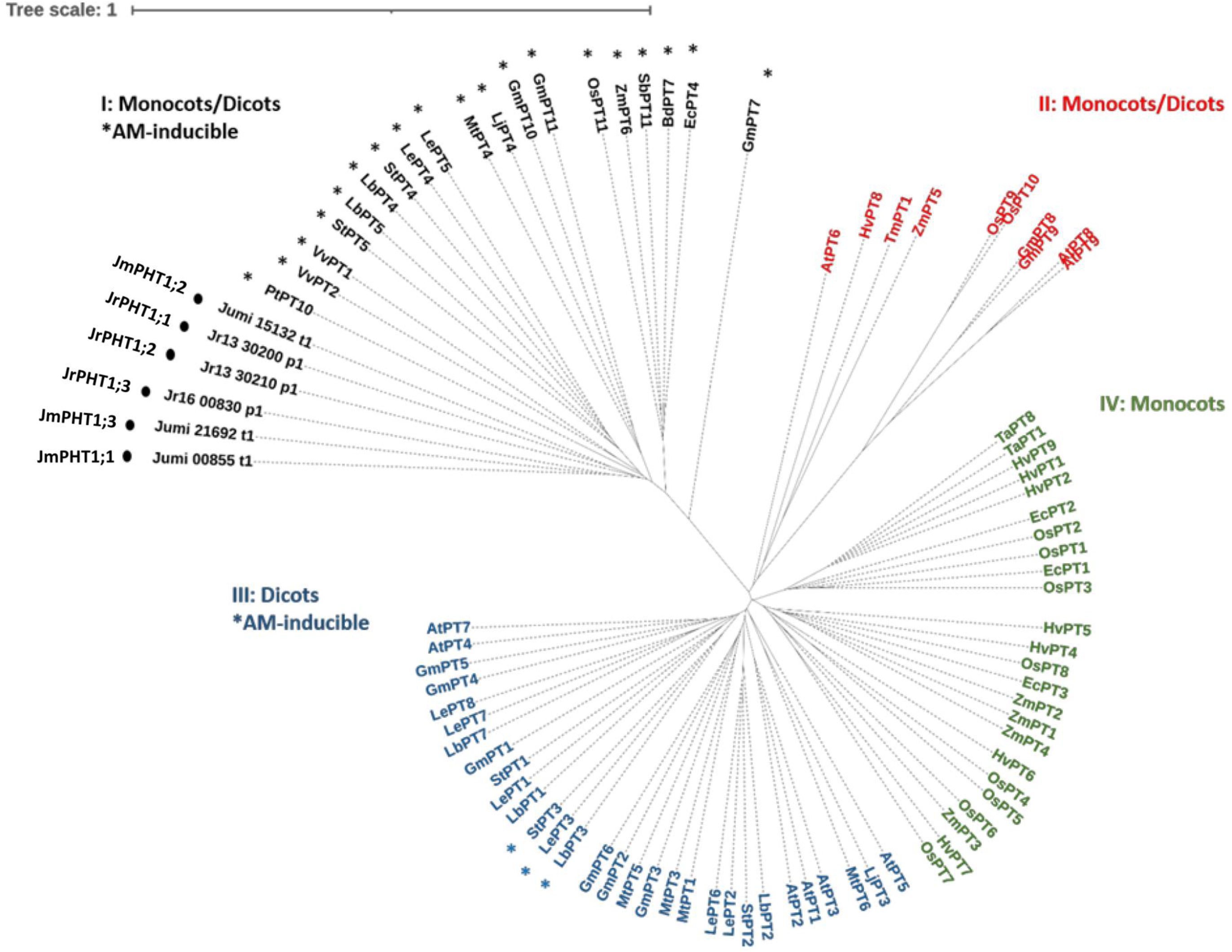
Unrooted phylogenic tree for the amino acid sequences of plant PHT1 transporters. Accessions numbers for protein sequences are detailed in Table S3. Roman numerical indicate the four PHT1 subfamilies thought to differ in evolutionary age (Nagy et al. 2005). Asterisks refer to AM-inducible *PHT1* genes (Table S3). Circles indicate the putative PHT1 transporters of *J. microcarpa* and *J. regia* orthologous to MtPT4. Abbreviations for plant species: At, *Arabidopsis thaliana*; Bd, *Brachypodium distachyon;* Ec, *Eleusine coracana*; Gm: *Glycine max*; Ha, Hv*, Hordeum vulgare*; Jm, *J. microcarpa;* Jr, *J.regia*; Lb, *Lycium barbarum*; Le, *Lycopersicon esculentum*; Lj, *Lotus japonicus*; Mt, *Medicago truncatula*; Os, *Oryza sativa*; Pt, *Populus trichocarpa;* Sb, *Sorghum bicolor*; St, *Solanum tuberosum*; Ta, *Triticum aestivum*; Tm, *Triticum monococcum;* Vv, *Vitis vinifera*; and Zm, *Zea mays*.

### 3.6 Phosphate transporter gene expression

With respect to plant Pi acquisition through the mycorrhizal pathway, we analysed the expression of the maize AM-essential phosphate transporter *ZmPHT1;6* owing to its unique identification through Orthofinder. As expected, analysis of transcript levels in non-mycorrhizal (NM_Jug) maize roots showed no expression of *ZmPHT1;6* **(Fig. 6**). By contrast, inoculation of the donor walnut RX1 (AM_Jug) with *R. irregularis* resulted in a significant (*p* < 0.05) induction of *ZmPHT1;6* in recipient maize roots whatever the Pi fertilization provided to maize. These results showed that in our experimental conditions, the expression of the gene *ZmPHT1;6* is induced by the colonization of maize roots through the formation of a common mycelial network irrespective of the Pi supply.

**Figure 6.**
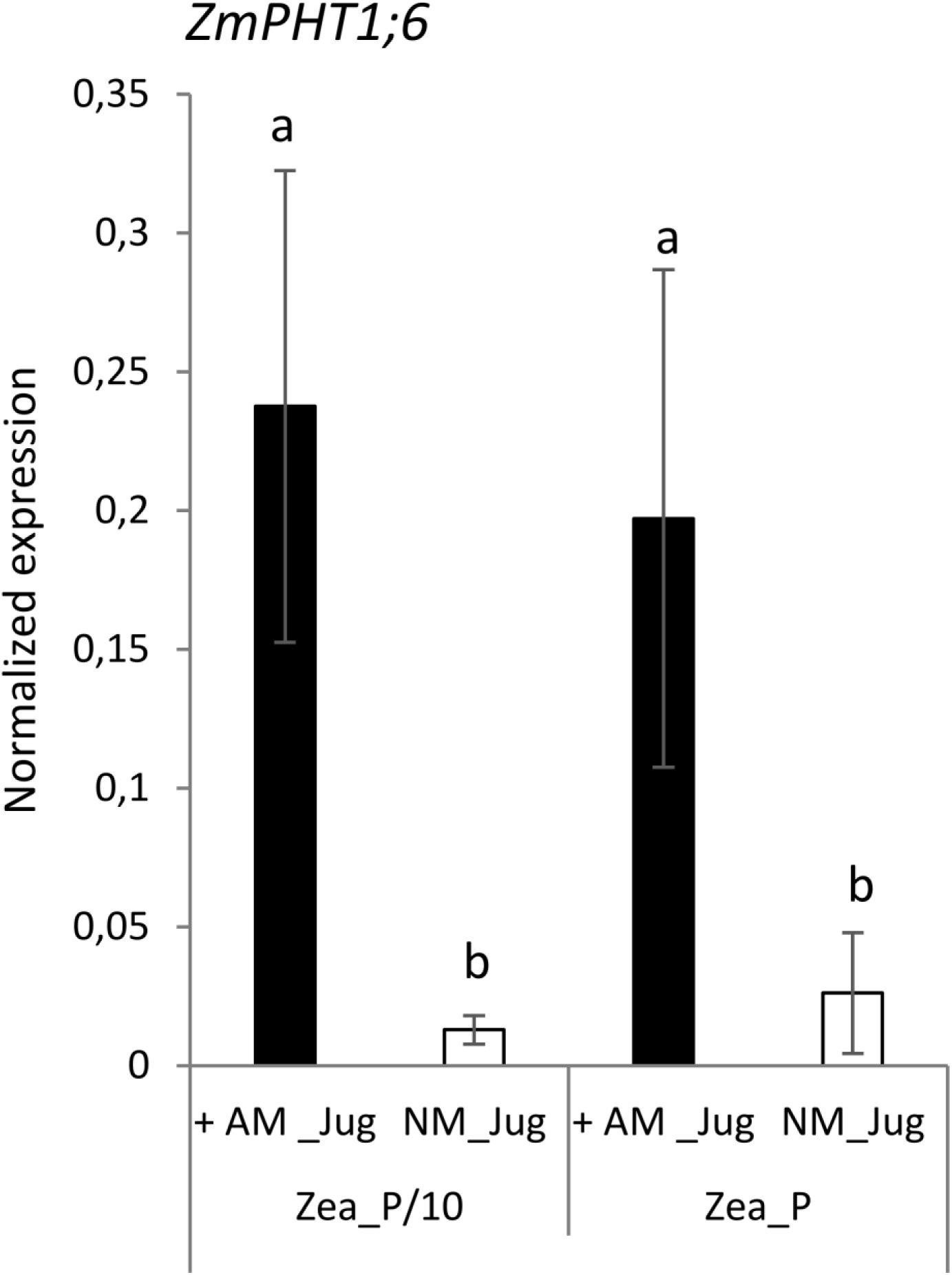
Expression pattern of the gene *ZmPHT1;6* coding the phosphate transporter orthologous to MtPT4 in recipient maize roots two months after the inoculation (AM_Jug) of donor walnut RX1 with *R. irregularis* rootstocks or not (NM_Jug) under contrasting Pi fertilization regimes of maize (Zea_P/10, Zea_P). Gene expression was quantified by RT-qPCR and expressed as normalized expression to the maize reference genes described in Table S1. Values correspond to the mean (±SE) of five replicates per treatment. Lower case letters indicate significant difference (*p* < 0.05) according to the Kruskal-Wallis H-test with post-hoc Tukey HSD.

Whatever the treatment of walnut donor plants, the results displayed in **Fig. S2** shows a very low expression in the roots of the hybrid rootstock RX1 of each of the three putative *J. microcarpa* Pi transporters retrieved in the MtPT4 orthogroup (JmPHT1;1, JmPHT1;2, JmPHT1;3). The analysis of the transcript abundance of *JrPHT1;1*; *JrPHT1;2*, and *JrPHT1;3* of *J. regia* in the roots of the hybrid walnut RX1 showed only weak expression levels of *JrPHT1;3* in the four conditions tested (**Fig.7)**. On the opposite, inoculation of RX1 (AM_Jug) with *R. irregularis* resulted in a significant (*p* < 0.05) induction of *JrPHT1;1* and *JrPHT1;2* in AM-walnut roots, relative to NM_Jug controls whatever the Pi fertilization provided to maize plants (**Fig. 7**). Expression levels of both *JrPHT1;1* and *JrPHT1;2* tend to be higher, although not significantly, under the P/10 than under the P supply to maize roots. Together these results demonstrated for the first time that the expression of *JrPHT1;1* and *JrPHT1;2* are induced in the roots of the hybrid walnut rootstock RX1 by *R. irregularis* irrespective of the Pi level supplied to maize recipient plants.

**Figure 7.**
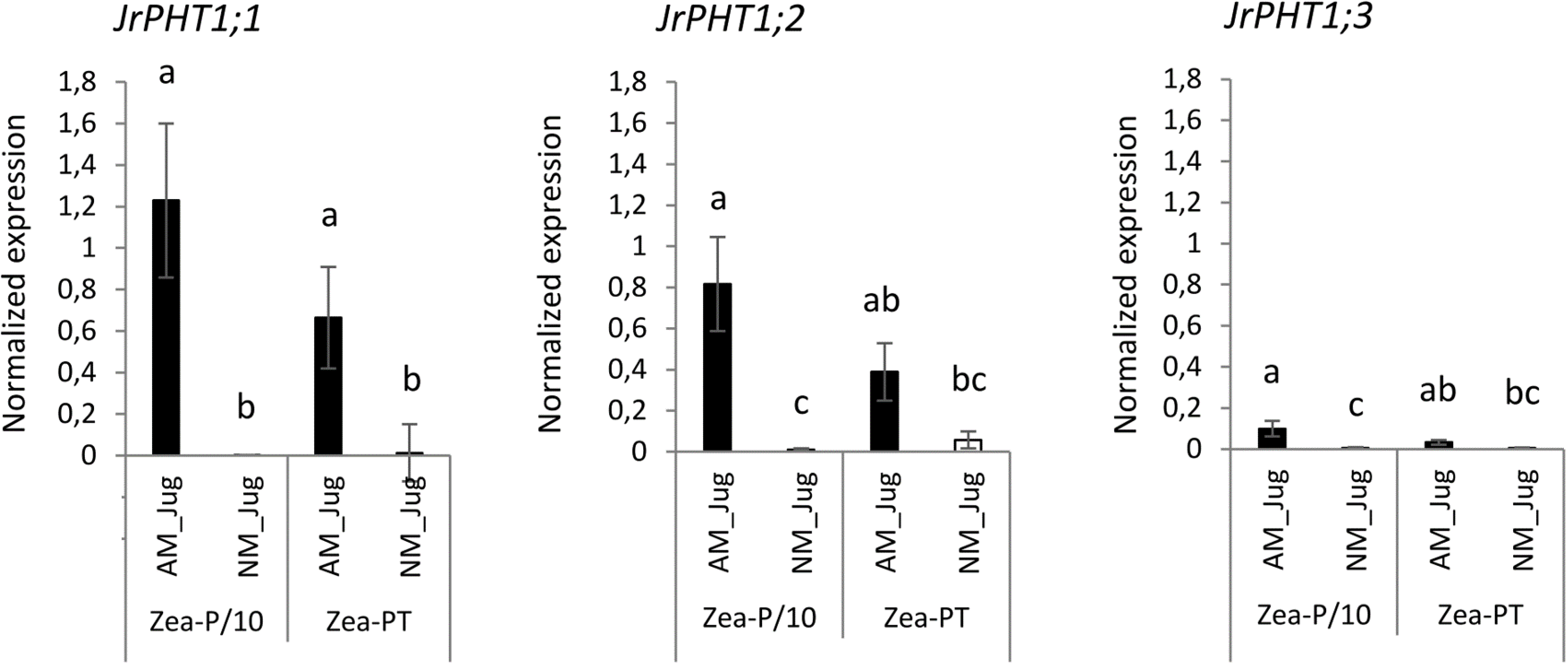
Expression patterns of the genes *JrPHT1;1*; *JrPHT1;2*, *JrPHT1*;3 coding proteins orthologous to the phosphate transporter MtPT4 in donor walnut RX1 roots two months after their inoculation with *R. irregularis* (AM_Jug) or not (NM_Jug) under contrasting Pi fertilization regimes of maize (Zea_P/10, Zea_P). Gene expression was quantified by RT-qPCR and expressed as normalized expression to the walnut reference genes described in Table S1. Values correspond to the mean (±SE) of five replicates per treatment. Lower case letters indicate significant difference (*p* < 0.05) according to the Kruskal-Wallis H-test with post-hoc Tukey HSD.

To monitor soil Pi acquisition through the fungal side, we monitored in walnut RX1 and maize roots the expression of the gene coding the high affinity phosphate transporter RiPT1 of *R. irregularis*. As expected, analysis of transcript levels in non-inoculated (NM_Jug) walnut and maize roots showed very low expression of *RiPT1* **(Fig. 8A, B**). By contrast, inoculation of RX1 (AM_Jug) with *R. irregularis* resulted in a significant (*p* < 0.05) induction of the expression of *RiPT1* not only in walnut **(Fig. 8A**), but also in maize roots **(Fig. 8B**), whatever the Pi fertilization provided to maize. These results indicated that the expression of *RiPT1* is induced in the mycorrhizated donor RX1 and but also in the recipient maize roots. Further Welch’s t-test comparison between AM walnut and AM maize roots, indicated a higher expression of the *RiPT1* gene in maize than in walnut roots both under the lowest (Zea_P/10, *p* < 0.08) and the highest Pi supply (Zea_P, *p* < 0.03). This showed that the contribution of the extra-radical mycelium of AM maize roots to Pi acquisition is higher than that of AM walnut roots.

**Figure 8.**
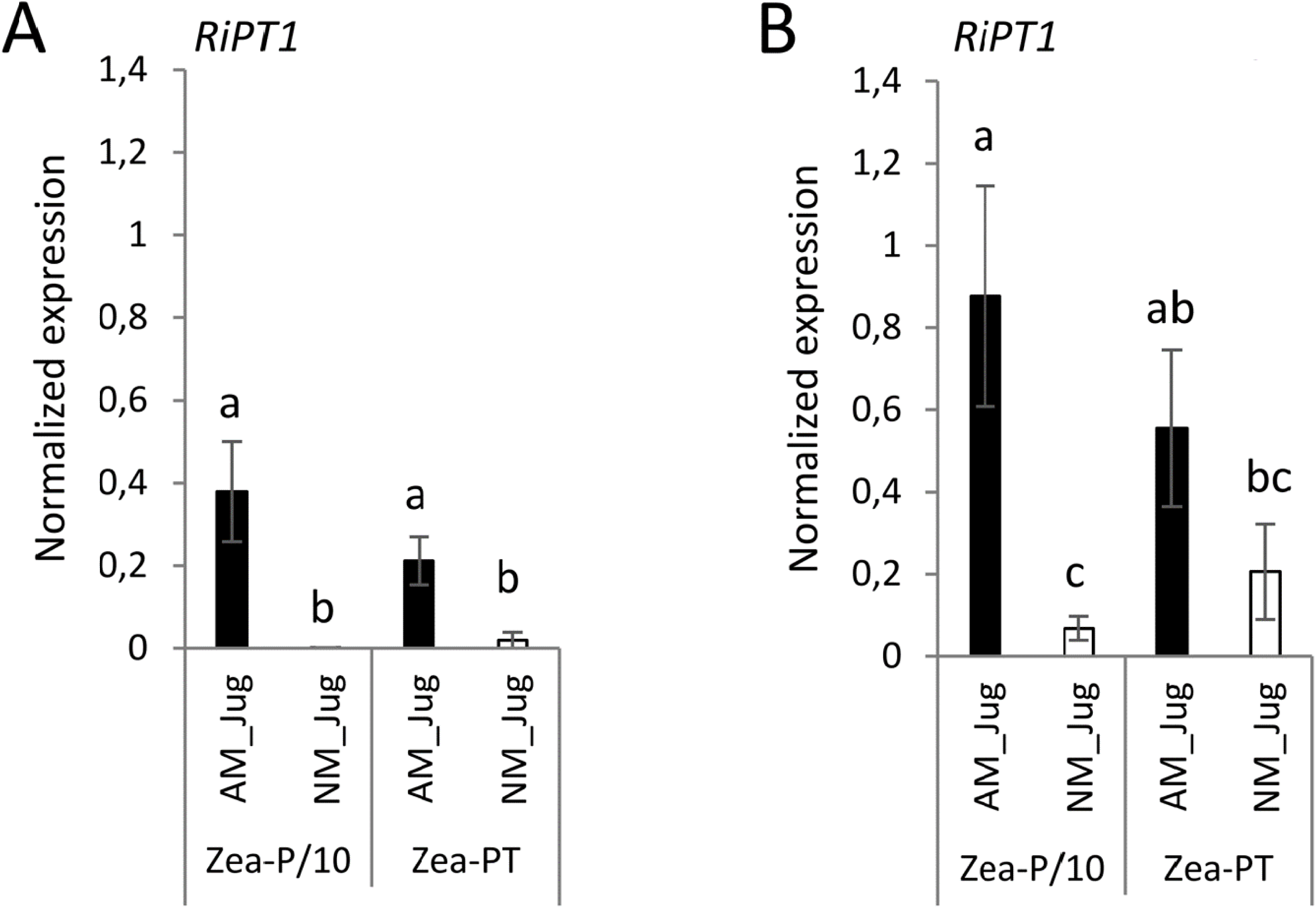
Expression pattern of the gene *RiPT1* of *R. irregularis* two months after its inoculation (AM_Jug) as compared to not inoculated (NM_Jug) of walnut RX1 rootstocks under contrasting Pi fertilization regimes of maize (Zea_P/10, Zea_P) in walnut donor (**A**), and non-inoculated maize recipient roots (**B**). Gene expression was quantified by RT-qPCR and expressed as normalized expression to the two fungal reference genes described in Table S1. Values correspond to the mean (±SE) of five replicates per treatment. Lower case letters indicate significant difference (*p* < 0.05) according to the Kruskal-Wallis H-test with post-hoc Tukey HSD.

## 4. Discussion

### 4.1 The inoculum reservoir formed by walnut roots allows the mycorrhization of maize through the formation of a CMN that gives access to maize fertilization residues

Irrespective of the Pi supplied to *Z. mays* recipient plants, non-inoculated maize roots connected to mycorrhizal walnut rootstocks displayed F, M, and A% values reaching 100, 30 and 25%, respectively. These results demonstrated that whatever the Pi fertilization regime applied to maize, the inoculum reservoir formed by AM walnut roots enables the mycorrhization of maize recipient roots through the development of a common mycelial network (CMN). Consistently, extra-radical hyphae emanating from walnut root were visualized in the two Pi treatments after trypan blue staining of the nylon mesh and substrate contained in the fungal pipes close to *Z. mays* roots. This observation showed that the extra-radical mycelium that developed outside AM walnut roots have crossed the 40-µm nylon mesh to reach the recipient compartment and further colonize maize roots. Consistently, the high affinity phosphate transporter *RiPT1* of *R. irregularis* and the maize phosphate transporter *ZmPHT1;6* were only expressed in maize connected to mycorrhizal RX1 donor plants.

There was also evidence that AM walnut plants connected to recipient roots have access to maize Pi fertilization residues via the CMN as shown by higher stem diameter, shoot and root dry biomasses recorded in mycorrhizal RX1 saplings as compared to non-inoculated walnuts in both Pi treatments. This finding demonstrated that walnut developing AM symbiosis gains growth benefit from the Pi supplied to maize through hyphal connections with walnut sapling. In addition, AM walnuts connected to recipient plants happened to be responsive to the extent of maize fertilization, as inferred from significantly lower M and A% in the P condition relative to the P/10 treatment. This showed that Pi shortage in the maize compartment sustains walnut mycorrhization, while a ten-fold higher Pi fertilization restricts the intra-radical development of *R. irregularis* in RX1. Earlier studies have reported a significant suppression of AM symbiosis upon high Pi supplies (Rausch et al. 2001; Breuillin et al. 2010; Bonneau et al. 2013; Kobae et al. 2016). When plants consume more Pi to provide photosynthetically fixed carbon than that furnished by mycorrhizal hyphae, the host limits soil Pi uptake through the mycorrhizal pathway by reducing carbon allocation to the symbiont in order to save energy (Treseder and Allen 2002; Blanke et al. 2005; Gavito et al. 2019). In agreement with this line, a decreased root carbon content, along with an increased leaf soluble Pi concentration and effective photochemical quantum yield of PS_II_ in AM walnuts compared to NM plants, was only significant in the Pi-limiting condition. In the process of phosphate suppression of mycorrhization, Pi acts systemically to repress the expression of symbiotic genes, including phosphate transporters (Breuillin et al. 2010), resulting in lower AMF colonization (Salmeron-Santiago et al. 2022). Consistently, the mean expression of *JrPHT1;1* and *JrPHT1;2* tend to be higher in the roots of RX1under the P/10 than under the P supply to maize roots. We concluded that in the walnut hybrid RX1, the AM fungus *R. irregularis* acts as a stronger carbon sink under Pi starvation (P/10) than under the Pi sufficient condition.

### 4.2 Common arbuscular mycorrhizal network alleviates Pi shortage in walnut and maize plants

Relative to a 1.3 mM Pi supply, a 0.13 mM Pi fertilization of non-mycorrhizal maize plants led to a significant reduction in stem height and diameter, leaf number, Pi and N content, shoot and root biomass. As previously observed (Almagrabi and Abdelmoneim 2012; Ma et al. 2021), this result showed that maize growth is limited by phosphate deficiency. In NM maize exposed to P sufficient treatments relative to Pi limiting conditions, higher leaf count, dry mass, plant height and stem thickness have also been reported: as an indicator of crop growth and development, when N is amply supplied, plant height is known to increase as P availability increases, similarly to stem thickness (Coetzee et al. 2016). In contrast to what observed in NM controls, in AM maize plants cultivated under the P/10 supply, all growth and nutritional parameters rose to a level similar to that recorded for NM plants under the P sufficient treatment, indicating that mycorrhization of maize plants alleviated Pi deficiency under the P/10 supply. These responses to mycorrhization in low P soil are consistent with previous studies demonstrating that maize crop benefits from mycorrhizal associations (Calderón-Vásquez et al. 2011; Ma et al. 2021). They also fit with the ‘optimal trade principle’ predicting that plant growth will most strongly benefit from symbiosis with AM fungi under conditions of low soil phosphorus supply (Johnson 2010; Johnson et al. 2015). Within this line, mean expression of the gene *RiPT1* in maize was higher under the P/10 than under the P supply. We concluded that under maize Pi shortage, the mycorrhizal connection between walnut and maize roots alleviates Pi deficiency in the recipient plant through a better mineral nutrition. This resulted in maize growth and nutritional benefits comparable to the P sufficient condition without a common arbuscular mycorrhizal network.

As expected from the absence of mycorrhiza connections between plants when walnut donor roots were not inoculated with *R. irregularis,* the Pi fertilization regime of maize had no impact on the parameters measured in NM walnut plants. This result supported the absence of solutes diffusion through the air gap. With regard to walnut growth benefits gained from the Pi supplied to maize through hyphal connections relative to non-fertilized NM walnut plants, we observed that a 0.13 mM Pi fertilization led to a significant increase in root collar, stem height, leaf Pi content, shoot and root DW. This underlines that under maize low Pi fertilization, the mycorrhizal connection between walnut and maize roots alleviates Pi deficiency in the donor plant through a better mineral nutrition. Contrary to maize, mycorrhizal growth benefits of walnut plants in terms of root collar and dry mass were conserved under high Pi fertilization of maize, even though intra-radical colonization was reduced. Altogether, these results support the idea that walnut is more dependent on the Pi uptake mycorrhizal pathway than maize to sustain its development owing to a coarse root architecture that has a limited intrinsic ability to absorb soil nutrients (Bates and Lynch 2001). We concluded that a common arbuscular mycorrhizal network is able to overcome phosphate shortage in walnut and maize plants.

### 4.3 *R. irregularis* acts as a stronger carbon sink in walnut than in maize

When comparing mycorrhizal walnut and maize plants grown under the P/10 supply relative to NM plants, a significant decreased root carbon content along with an increased effective photochemical quantum yield of PS_II,_ was only observed in RX1, while both plants displayed a higher leaf Pi concentration. When assessing the S/R biomass ratio as a measure of the carbon cost for AM establishment (Chen et al. 2010), mycorrhizal colonization decreased the shoot to root biomass partitioning only in RX1 grown under the P/10 supply. These findings indicated that AM fungal colonization enhances walnut investment in root biomass development. Taken together, these results showed that mycorrhization-induced Pi uptake generates a higher carbon cost for walnut donor plants than for maize plants by increasing walnut plant photosynthesis to feed the AM fungus with carbon assimilate. In support for this rationale, when using walnut trees (*Juglans nigra* L., C3-plant) and maize (C4-plant), which display distinctly different ratios of ^13^C/^12^C (∂^13^C), analysis of ∂ ^13^C from an AM mycelium taken from the surrounding environment of intercropped walnut and maize roots indicated the transfer of walnut photosyntates to the AM mycelium (van Tuinen et al. 2020). With regard to the potential mechanisms underlying a larger carbon investment in AM symbiosis of walnut than maize, Graham and Esseinstat (1994) proposed that a high mycorrhizal dependency, as displayed by walnut, may reflect a greater availability of the carbon supplied by the host. On the opposite, because plants with rapid growth rate need more carbon and absorb more soil nutrients to achieve fast growth (Lovelock 2008), less photosynthates may thus be available to feed the CMN. According to Elser et al. (2000), plant growth rate is usually negatively related to C/N content ratio. In the current study, mycorrhization of maize with *R. irregularis* through the CMN led when compared to NM plants, to a significant decrease in leaf C/N ratio under the maize Pi-limited condition, but not in the Pi-replete condition. As this result was not observed in walnut rootstocks, our data support the idea that at low soil fertility, walnut plant growth is limited more by nutrients uptake than by carbon supply (Poorter and De Jong 1999; Korner 2003). Under these conditions, it is more advantageous for walnut saplings than for maize plant to allocate carbon assimilates to feed the AM symbiont (van der Heijden and Horton 2009).

## 5. Conclusion

In agreement with our working hypotheses, this study provides for the first time evidence that (1) the inoculum reservoir formed by walnut roots allows the mycorrhization of maize roots through the formation of a CMN, thereby giving access to maize fertilization residues; (2) hyphal connections under shortage of P result in a benefit comparable to P sufficient condition without a CMN; and (3) due to a slower root growth rate than maize, walnut invests more carbon in the development of the AM fungus than maize does. These findings support the idea that in an agroforestry system the presence of perennial root systems enable the maintenance of an active AM fungal network, which constitutes an AM reservoir allowing root colonization of the annual crop, and authorizing low phosphorus fertilization input. CMN formation is a facilitative mechanisms between trees and companion crops favourable to their growth and development (Bainard et al. 2011; Bainard et al. 2012; Ingleby et al. 2007). These results are also in accordance with the stress-gradient hypothesis that predicts that low-fertilised agricultural lands would benefit more from intercropping than high-fertilised lands (Brooker and Callaghan 1998; Zhu et al. 2023). Therefore, mycorrhization of tree saplings with a suitable AM fungus able to develop CMN favourable to alley annual crop may result in a benefit comparable to P fertilization and would therefore be a recommendable agronomical practice in the framework of the agroecological transition of agricultural systems.

## Acknowledgements

Emma Mortier’s PhD thesis was funded by The French Ministry of Agriculture and benefited from a collaboration with L Jouve’s company. We would like to thank all the people who provided us with invaluable assistance in carrying out this work.

## Contributions

Conceptualization, E.M., F.M-L, G.R and O.L.; methodology, E.M, A.M; J.K, G.R, and O.L. writing—original draft, —review and editing: E.M, A.M; J.K, F.M-L, G.R, and O.L.; funding acquisition, F. M-L.; supervision, F. M-L, G.R and O.L.

## Ethics declarations

Ethics approval : Not applicable.

Consent to participate : Not applicable.

Consent for publication : Not applicable.

Conflict of interest : The authors declare no competing interests.

## Code availability

Not applicable.

## Data availability

The datasets generated during and/or analyzed during the current study are available from the corresponding author on reasonable request.

## Figure Legends

**Figure S1.** Sequence alignment of the mycorrhiza-inducible phosphate transporter *M. truncatula* PT4 (Medtr1g028600.1) and its corresponding orthologs in *J. regia* (JrPHT1;1/Jr13_30200_p1, JrPHT1;2/Jr13_30210_p1; JrPHT1;3/Jr16_00830_p1), *J. microcarpa* (JmPHT1;1/ jumi_00855.t1, JmPHT1;2/jumi_15132.t1; JmPHT1;3/jumi_21692.t1), and *Z. mays* (Zm00001d011498_P001). Protein sequences were aligned using MultAlin (http://multalin.toulouse.inra.fr/multalin/). Green bars indicate predicted transmembrane segments according to DeepTMHMM (https://dtu.biolib.com/DeepTMHMM). Yellow and red bar underline the sequence signature GGDYPLSATIxSE and the intracellular central loop conserved between PHT1 proteins, respectively. Cytoplasmic (In) and exoplasmic (Out) sequence orientations were predicted according to DeepTMHMM (https://dtu.biolib.com/DeepTMHMM).

**Figure S2.** Expression patterns of the genes *JmPHT1;1*; *JmPHT1;2*, *JmPHT1*;3 coding proteins orthologous to the phosphate transporter MtPT4 in donor walnut RX1 roots two months after their inoculation with *R. irregularis* (AM_Jug) or not (NM_Jug) under contrasting Pi fertilization regimes of maize (Zea_P/10, Zea_P). Gene expression was quantified by RT-qPCR and expressed as normalized expression to the walnut reference genes described in Table S1. Values correspond to the mean (±SE) of five replicates per treatment. Lower case letters indicate significant difference (*p* < 0.05) according to the Kruskal-Wallis H-test with post-hoc Tukey HSD.

## Notes

### Competing Interest Statement

The authors have declared no competing interest.

