## Supplemental Figures and Tables for "Evidence that common arbuscular mycorrhizal network alleviates phosphate shortage in interconnected walnut sapling and maize plants"

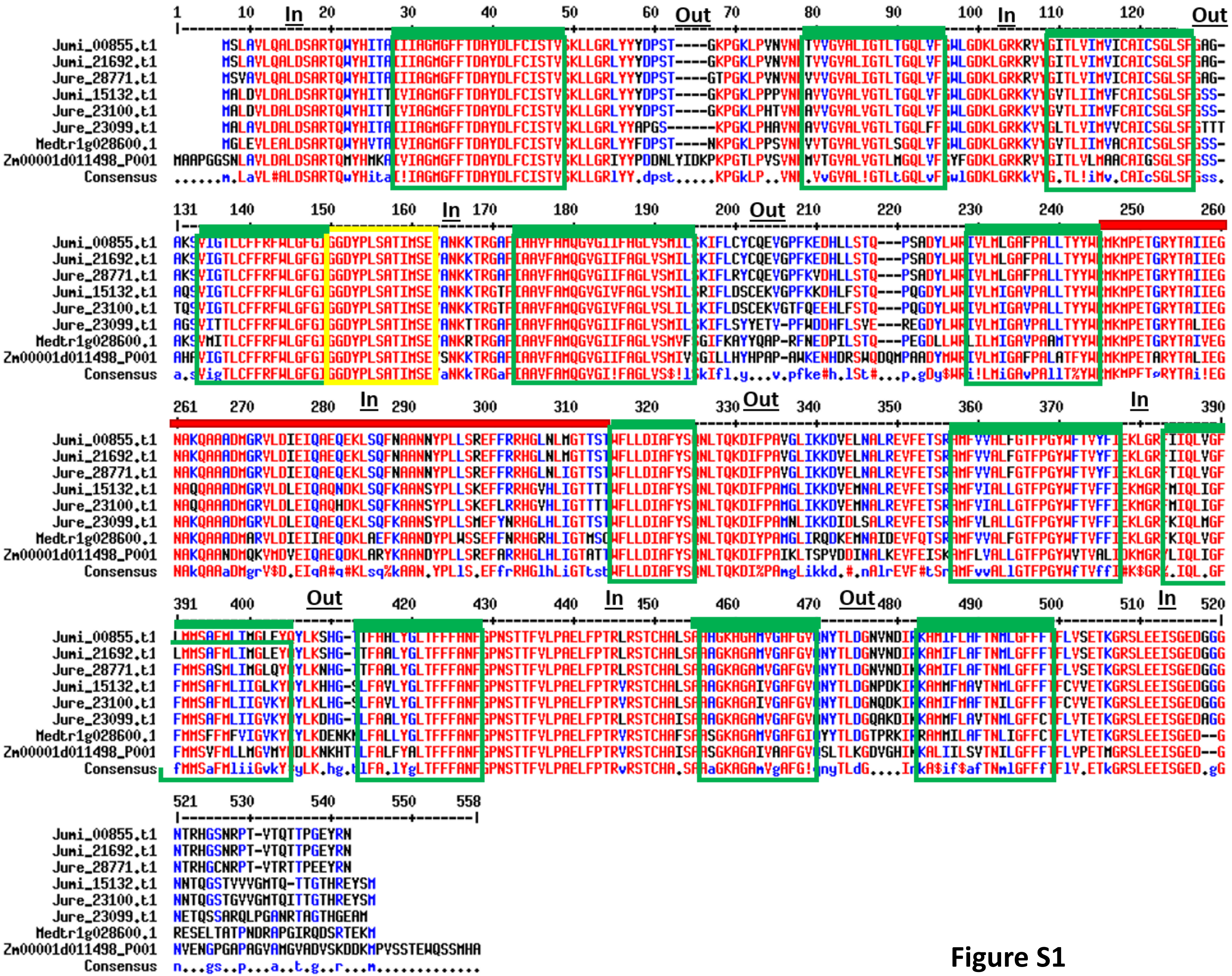
In
Out
Jumi_00855.t1
Jumi_21692.t1
Jure_28771.t1
Jumi_15132.t1
Jure_23100.t1
Jure_23099.t1
Medtr1g028600.1
Zm00001d011498_P001
Consensus

**Figure S1**

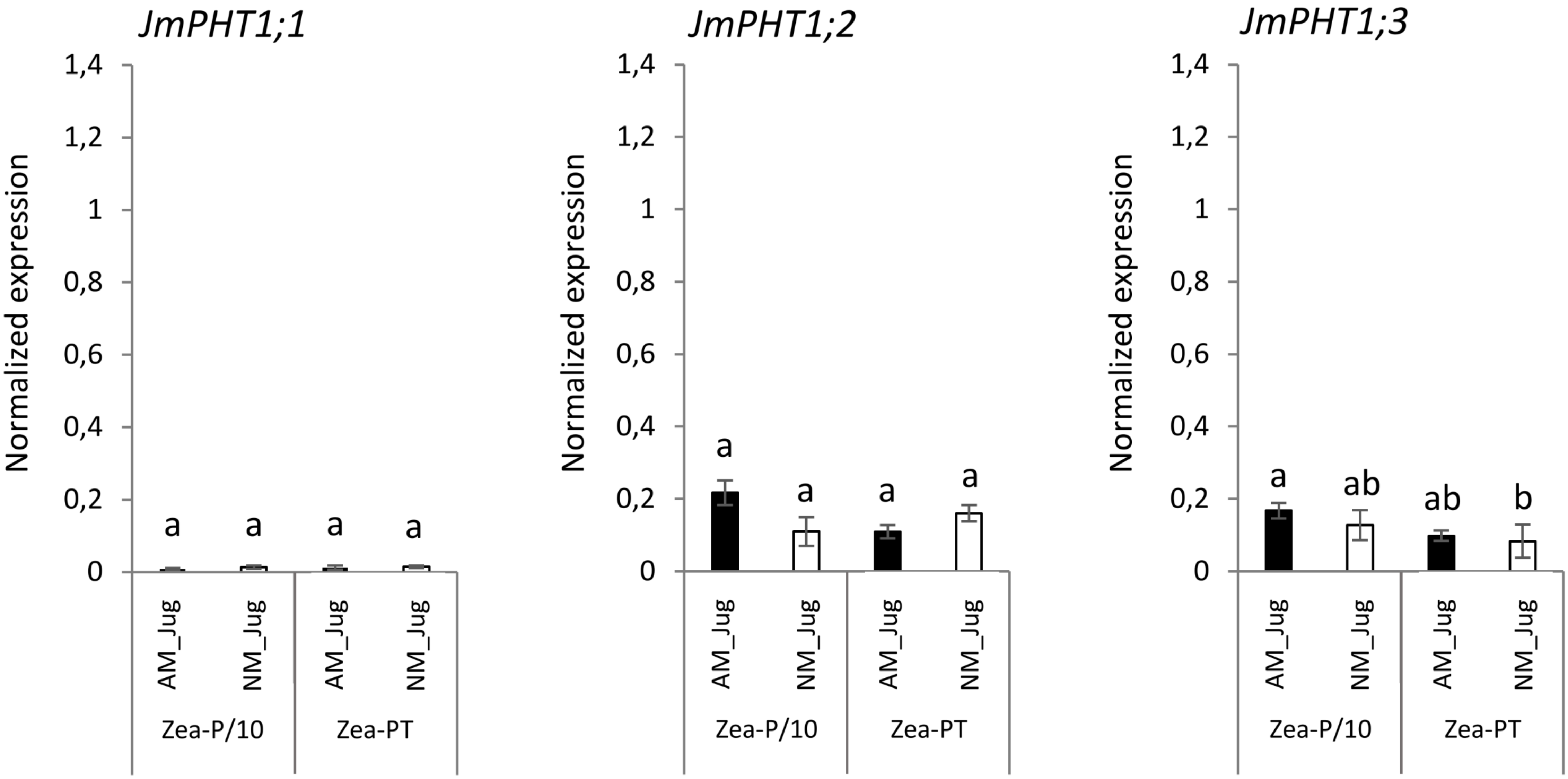
JmPHT1;1
JmPHT1;2
JmPHT1;3
Normalized expression
1,4
1,2
1
0,8
0,6
0,4
0,2
0
a
ab
b
AM_Jug
NM_Jug
Zea-P/10
Zea-PT

**Figure S2**

### SUPPLEMENTARY TABLES

**Table S1.** Primers used for qRT-PCR in *J. regia*, *J. microcarpa*, *Z. mais*, and *R. irregularis*

| **Oligoname** | **Sequence** | **Transcript ID/Reference** |
|---|---|---|
| ***J. regia*** | | |
| *JregGAPDHrefF* | ATCATGGGTAAAGACCCCGC | XM_018984056.1/ Zhou et al. (2018) |
| *JregGAPDHrefR* | TCAGTGACCGCATCCTTAGC | |
| *JregACT2refF* | ATGGTCCCAAACATGACCCA | XM_018959949.1/Zhou et al. (2018) |
| *JregACT2refR* | TGTTGGAGGAGCTTGTGCAG | |
| *JregPT4-28771F* | TCGAGAAGCTGGGTCGTTTC | Jr16_00830_p1/This study |
| *JregPT4-28771R* | TTCAGGAGTGGTCCGAGTCA | |
| *JregPT4-1F* | GTTGCCCTTGTTGGTACCCT | Jr13_30200_p1/This study |
| *JregPT4-23100R* | AGCATATGGCGCAAAACACC | |
| *JregPT4-23099F* | CCTGGCAGTAAGCCTGGAAA | Jr13_30210_p1 /This study |
| *JregPT4-23099R* | AGTTTATCACCCAGCCAGCC | |
| ***J. microcarpa*** | | |
| *JmicGAPDHrefF* | CCCTCACAGCGAAACGACC | Jumi_01067.t1/This study |
| *JmicGAPDHrefR* | TTTTTGAAGAATGGTGATGCAGGC | |
| *JmicACT2refF* | ACGGGTGGGGAAGATTTGAC | Jumi_26547.t1/This study |
| *JmicACT2refR* | GGAGGGAGCACCATGTATCC | |
| *JmicPT4-2192F* | ATGTCCCCGAACTCCTCTATCTT | Jumi_21692.t1/This study |
| *JmicPT4-2192R* | CGATTGTGTTGTCGCAGGG | |
| *JmicPT4-15132F* | GCCCAAGACTCCTTAACCCC | Jumi_15132.t1/This study |
| *JmicPT4-15132R* | CATCGACGAGGCTGATACGA | |
| *JmicPT4-855F* | AGAACCAGAGAACAGTGGCG | Jumi_00855.t1/This study |
| *JmicPT4-855R* | CATGGACCCTCACCTTCGAC | |
| ***Z. mays*** | | |
| *Zm B-tubulinrefF* | CTACCTCACGGCATCTGCTATGT | Lin et al. (2014) |
| *Zm B-tubulinrefR* | GTCACACACACTCGACTTCACG | |
| *Zm-EF1*αrefF | TGGGCCTACTGGTCTTACTACTGA | Lin et al. (2014) |
| *Zm-EF1αrefF* | ACATACCCACGCTTCAGATCCT | |
| *ZmPT1;6 F* | TTCTGCATCTCCACCGTGTC | Zm00001d011498_P001= ZmPht1;6 / Glassop et al. (2005) |
| *ZmPT1;6 R* | CGCCTGTCACCATGTTGTTG | |
| ***R. irregularis*** | | |
| *Riα-tubulin a1 F* | TGTCCAACCGGTTTTAAAGT | TC105406/Gomez et al. (2009) |
| *Riα-tubulin a1 R* | AAAGCACGTTTGGCGTACAT | |
| *RiTEFrefF* | TGACAGGCGATCTGGTAAGG | Calabrese et al. (2019) |
| *RiTEFrefR* | TCAGCGAAGGTCTCAACCAC | |

| | | |
|---|---|---|
| *RiPT1F* | TGCTCGGTTGTGGTGCTATT | TC345640/Calabrese et al. (2019) |
| *RiPT1R* | CGACGTCCGAAGTTGCTTTG | |

**Table S2.** Effect of AM colonization and Pi availability on plant growth and nutritional parameters. Values correspond to the mean (±SE) of six replicates per treatment. Lower case letters indicate significant difference (Kruskal-Wallis H-test with post-hoc Tukey HSD; $p < 0.05$).

| | Zea_P/10 | | Zea_P | | |
|---|---|---|---|---|---|
| ***Walnut*** | M | NM | M | NM | pvalue |
| Root length (cm) | 33.13±5.1 | 46.42±4.1 | 38.10±4.3 | 38.73±5.2 | 0.47 |
| **Collar (mm)** | **3.78±0.3** $^{ab}$ | **2.93±0.2** $^{c}$ | **3.99±0.2** $^{a}$ | **3.04±0.2** $^{bc}$ | **<0.01** |
| **Height (cm)** | **17.02±1.7** $^{a}$ | **10.10±1.9** $^{b}$ | **12.84±0.8** $^{ab}$ | **10.10±1.2** $^{b}$ | **0.02** |
| Leaf number | 5.17±0.7 | 4.50±0.5 | 4.29±0.4 | 5.14±0.5 | 0.56 |
| **Shoot dw (g)** | **1.26±0.2** $^{a}$ | **0.51±0.1** $^{b}$ | **1.06±0.1** $^{a}$ | **0.57±0.1** $^{b}$ | **<0.01** |
| **Root dw (g)** | **2.50±0.3** $^{a}$ | **0.81±0.2** $^{b}$ | **1.85±0.2** $^{a}$ | **0.94±0.3** $^{b}$ | **<0.01** |
| **Shoot/root** | **0.49±0.03** $^{b}$ | **0.86±0.2** $^{a}$ | **0.58±0.03** $^{ab}$ | **0.68±0.1** $^{a}$ | **0.05** |
| Fv_fm | 0.72±0.02 | 0.71±0.02 | 0.70±0.01 | 0.71±0.01 | 0.90 |
| **$Y_{II}$** | **0.45±0.01** $^{a}$ | **0.34±0.02** $^{b}$ | **0.44±0.01** $^{a}$ | **0.42±0.03** $^{a}$ | **<0.01** |
| Root N (%DW) | 1.5±0.1 | 1.6±0.1 | 1.7±0.1 | 1.7±0.1 | 0.33 |
| **Root C (%DW)** | **39.6±0.3** $^{b}$ | **42.8±0.3** $^{a}$ | **39.9±1.4** $^{b}$ | **40.3±0.1** $^{b}$ | **0.02** |
| Root P (nmole/mg) | 40.18±4.2 | 45.97±6.3 | 50.56±2.6 $^{a}$ | 41.82±3.1 $^{a}$ | 0.13 |
| Leaf N (% DW) | 2.8±0.1 | 2.7±0.1 $^{a}$ | 2.6±0.04 $^{a}$ | 2.8±0.1 $^{a}$ | 0.18 |
| Leaf C (% DW) | **45.4±0.3** $^{a}$ | **44.3±0.4** $^{ab}$ | **44.5±0.1** $^{ab}$ | **43.8±0.4** $^{b}$ | **0.04** |
| **Leaf P (nmole/ mg)** | **62.0±1.6** $^{a}$ | **49.1±4.7** $^{b}$ | **53.3±4.5** $^{ab}$ | **65.9±5.4** $^{a}$ | **0.03** |
| Leaf C/N | 16.2±0.5 | 16.6±0.6 | 17.3±0.3 | 15.6±0.9 | 0.3 |

| | Zea_P/10 | | Zea_P | | |
|---|---|---|---|---|---|
| ***Maize*** | M | NM | M | NM | pvalue |
| Root length (cm) | 59.0±4.2 | 45.3±2.5 | 53.7±2.4 | 60.7±6.7 | 0.05 |
| **Collar (mm)** | **16.8±0.4** $^{a}$ | **12.3±0.9** $^{b}$ | **17.4±0.4** $^{a}$ | **16.8±0.8** $^{a}$ | **<0.01** |
| **Height (cm)** | **135.5±8.8** $^{a}$ | **82.6±10.8** $^{b}$ | **132.1±11.1** $^{a}$ | **129.9±12.3** $^{a}$ | **0.02** |
| Leaf number | **8.2±0.3** $^{a}$ | **6.6±0.3** $^{b}$ | **8.6±0.2** $^{a}$ | **8.4±0.2** $^{a}$ | **<0.01** |
| **Shoot DW (g)** | **3.6±0.7** $^{a}$ | **2.4±0.4** $^{b}$ | **6.8±1.0** $^{a}$ | **5.6±0.7** $^{a}$ | **<0.01** |
| **Root DW (g)** | **1.2±0.1** $^{a}$ | **0.4±0.1** $^{b}$ | **0.9±0.1** $^{a}$ | **0.9±0.1** $^{a}$ | **<0.01** |
| Shoot/root | **5.6±0.5** $^{b}$ | **5.5±0.5** $^{b}$ | **8.1±0.6** $^{a}$ | **6.7±0.7** $^{ab}$ | **0.04** |
| Fv_fm | **0.8±0.01** $^{ab}$ | **0.8±0.01** $^{a}$ | **0.8±0.01** $^{a}$ | **0.8±0.01** $^{a}$ | **<0.01** |
| $Y_{II}$ | 0.46±0.01 | 0.45±0.01 | 0.43±0.01 | 0.46±0.01 | 0.3 |
| Root N (% DW) | 1.7±0.2 | 1.7±0.1 | 1.5±0.2 | 1.7±0.1 | 1.0 |

| | | | | | |
|---|---|---|---|---|---|
| Root C (% DW) | 31.4±2.5 | 34.4±1.6 | 32.9±2.7 | 33.7±1.8 | 0.8 |
| Root Pi (nmole Pi/mg) | 66.3±7.7 | 42.9±1.9 | 52.4±3.3 | 47.6±8.1 | 0.1 |
| **Leaf N** (% DW) | **2.9±0.1** $^{a}$ | **2.6±0.1** $^{b}$ | **3.0±0.1** $^{a}$ | **2.9±0.2** $^{a}$ | **0.02** |
| Leaf C (% DW) | 44.1±0.3 | 44.1±0.1 | 44.3±0.2 | 44.7±0.2 | 0.1 |
| **Leaf Pi (nmole Pi/mg)** | **102.19±5.6** $^{a}$ | **79.3±2.6** $^{b}$ | **123.8±7.2** $^{a}$ | **115.7±11.4** $^{a}$ | **<0.01** |
| **Leaf C/N** | **15.1±0.5** $^{b}$ | **17.1±0.4** $^{a}$ | **14.8±0.4** $^{b}$ | **15.6±0.9** $^{ab}$ | **0.02** |

**Table S3**. Proteins of *J. regia*, *J. microcarpa* and *Z. mays* contained in the orthogroup of the mycorrhiza-inducible PHT1 transporter MtPT4 of *M. truncatula*.

| **Protein name** | **Organism** | **Locus/Accession** | **Source** |
|---|---|---|---|
| MtPT4 | *M. truncatula* | Medtr1g028600; Q8GSG4.1 | https://genome.jgi.doe.gov/portal/pages/dynamicOrganismDownload.jsf?organism=Mtruncatula/Mtruncatula_198_protein.fa.gz; https://www.uniprot.org/ |
| JrPHT1;1 | *J. regia* | Jr13_30200_p1; A0A2I4GU17 | https://treegenesdb.org/FTP/Genomes/Jure/v2.0/annotation/Jure.2_0.pep.fa.gz |
| JrPHT1;2 | *J. regia* | Jr13_30210_p1; A0A2I4GU05 | https://treegenesdb.org/FTP/Genomes/Jure/v2.0/annotation/Jure.2_0.pep.fa.gz |
| JrPHT1;3 | *J. regia* | jr16_00830_P1; A0A6P9EI50 | https://treegenesdb.org/FTP/Genomes/Jure/v2.0/annotation/Jure.2_0.pep.fa.gz |
| JmPHT1;1 | *J. microcarpa* | jumi_00855.t1 | https://treegenesdb.org/FTP/Genomes/Jumi/v1.0/annotation/jumi.1_0.peptides.fa |
| JmPHT1;2 | *J. microcarpa* | jumi_15132.t1 | https://treegenesdb.org/FTP/Genomes/Jumi/v1.0/annotation/jumi.1_0.peptides.fa |
| JmPHT1;3 | *J. microcarpa* | jumi_21692.t1 | https://treegenesdb.org/FTP/Genomes/Jumi/v1.0/annotation/ jumi.1_0.peptides.fa |
| Zm PHT1;6 | *Z. mays* | Zm00001d011498_P001/ NP_001105776.1 | https://ftp.ensemblgenomes.org/pub/release-43/plants/fasta/zea_mays/pep/Zea_mays.B73_RefGen_v4.pep.all.fa.gz; https://www.ncbi.nlm.nih.gov/ |

**Table S4**. List of the plant PHT1 transporters used to build the phylogenic tree.

| Protein name | Organism | Accession | Reference | DOI |
|---|---|---|---|---|
| AtPT1 | *Arabidopsis thaliana* | NP_199149.1 | Tabata et al. 2000 | https://doi.org/10.1038/35048507 |
| AtPT2 | *Arabidopsis thaliana* | NP_001190462.1 | Tabata et al. 2000 | https://doi.org/10.1038/35048507 |
| AtPT3 | *Arabidopsis thaliana* | NP_199150.1 | Tabata et al. 2000 | https://doi.org/10.1038/35048507 |
| AtPT4 | *Arabidopsis thaliana* | NP_181428.1 | Lin et al. 1999 | https://doi.org/10.1038/45471 |
| AtPT5 | *Arabidopsis thaliana* | NP_180842.1 | Lin et al. 1999 | https://doi.org/10.1038/45471 |
| AtPT6 | *Arabidopsis thaliana* | NP_199148.1 | Tabata et al. 2000 | https://doi.org/10.1038/35048507 |
| AtPT7 | *Arabidopsis thaliana* | NP_001319749.1 | Salanoubat et al, 2000 | https://doi.org/10.1038/35048706 |
| AtPT8 | *Arabidopsis thaliana* | NP_001323235.1 | Theologis et al. 2000 | https://doi.org/10.1038/35048500 |
| AtPT9 | *Arabidopsis thaliana* | NP_177769.1 | Theologis et al. 2000 | https://doi.org/10.1038/35048500 |
| BdPT7 | *Brachypodium distachyon* | XP_003569484.1 | Hong et a. 2002 | https://doi.org/10.1007/s00425-012-1677-z |
| EcPT1 | *Eleusine coracana* | AJD86034 | Pudake et al. 2017 | https://doi.org/10.1007/s13205-017-0609-9 |
| EcPT2 | *Eleusine coracana* | AJD86035.1 | Pudake et al. 2017 | https://doi.org/10.1007/s13205-017-0609-9 |
| EcPT3 | *Eleusine coracana* | AJD86036.1 | Pudake et al. 2017 | https://doi.org/10.1007/s13205-017-0609-9 |
| EcPT4 | *Eleusine coracana* | AJD86037.1 | Pudake et al. 2017 | https://doi.org/10.1007/s13205-017-0609-9 |
| GmPT1 | *Glycine max* | NP_001239971.1 | Fan et al. 2013 | https://doi.org/10.1186/1471-2229-13-48 |
| GmPT2 | *Glycine max* | NP_001240239.1 | Fan et al. 2013 | https://doi.org/10.1186/1471-2229-13-48 |
| GmPT3 | *Glycine max* | NP_001241164.1 | Fan et al. 2013 | https://doi.org/10.1186/1471-2229-13-48 |
| GmPT4 | *Glycine max* | NP_001304639.2 | Fan et al. 2013 | https://doi.org/10.1186/1471-2229-13-48 |
| GmPT5 | *Glycine max* | NP_001304588.2 | Wang et al. 2019 | https://doi.org/10.1186/s12870-019-1959-8 |
| GmPT6 | *Glycine max* | NP_001240032.1 | Fan et al. 2013 | https://doi.org/10.1186/1471-2229-13-48 |
| GmPT7 | *Glycine max* | NP_001239765.1 | Chen et al. 2019 | https://doi.org/10.1111/nph.15541 |
| GmPT8 | *Glycine max* | NP_001345390.1 | Fan et al. 2013 | https://doi.org/10.1186/1471-2229-13-48 |
| GmPT9 | *Glycine max* | NP_001241574.1 | Fan et al. 2013 | https://doi.org/10.1186/1471-2229-13-48 |
| GmPT10 | *Glycine max* | NP_001241400.1 | Fan et al. 2013 | https://doi.org/10.1186/1471-2229-13-48 |
| GmPT11 | *Glycine max* | NP_001241127.1 | Fan et al. 2013 | https://doi.org/10.1186/1471-2229-13-48 |
| HvPT1 | *Hordeum vulgare* | AAN37900.1 | Rae et al. 2003 | https://doi.org/10.1023/B:PLAN.0000009259.75314.15 |
| HvPT2 | *Hordeum vulgare* | AAO72434.1 | Smith et al. 1999 | https://doi.org/10.1007/978-94-017-2685-6_19 |
| HvPT4 | *Hordeum vulgare* | AAO72437.1 | Smith et al. 1999 | https://doi.org/10.1007/978-94-017-2685-6_19 |
| HvPT5 | *Hordeum vulgare* | AAO72435.1 | Smith et al. 1999 | https://doi.org/10.1007/978-94-017-2685-6_19 |
| HvPT6 | *Hordeum vulgare* | AAN37901.1 | Smith et al. 1999 | https://doi.org/10.1007/978-94-017-2685-6_19 |
| HvPT7 | *Hordeum vulgare* | AAO72436.1 | Smith et al. 1999 | https://doi.org/10.1007/978-94-017-2685-6_19 |
| HvPT8 | *Hordeum vulgare* | AAO72440.1 | Smith et al. 1999 | https://doi.org/10.1007/978-94-017-2685-6_19 |
| JmPHT1;1 | *Juglans microcarpa* | jumi_00855.t1 | this study | |
| JmPHT1;2 | *Juglans microcarpa* | jumi_15132.t1 | this study | |

| | | | | |
|---|---|---|---|---|
| JmPHT1;3 | *Juglans microcarpa* | jumi_21692.t1 | this study | |
| JrPHT1;1 | *Juglans regia* | Jr13_30200_p1, A0A2I4GU17 | this study | |
| JrPHT1;2 | *Juglans regia* | Jr13_30210_p1, A0A2I4GU05 | this study | |
| JrPHT1;3 | *Juglans regia* | Jr16_00830_p1, A0A6P9EI50 | this study | |
| LbPT1 | *Lycium barbarum* | AIU41746.1 | Hu et al. 2017 | https://doi.org/10.1093/treephys/tpw125 |
| LbPT2 | *Lycium barbarum* | AIU41747.1 | Hu et al. 2017 | https://doi.org/10.1093/treephys/tpw125 |
| LbPT3 | *Lycium barbarum* | AIU41748.1 | Hu et al. 2017 | https://doi.org/10.1093/treephys/tpw125 |
| LbPT4 | *Lycium barbarum* | AIU41749.1 | Hu et al. 2017 | https://doi.org/10.1093/treephys/tpw125 |
| LbPT5 | *Lycium barbarum* | AIU41750.1 | Hu et al. 2017 | https://doi.org/10.1093/treephys/tpw125 |
| LbPT7 | *Lycium barbarum* | AIU41751.1 | Hu et al. 2017 | https://doi.org/10.1093/treephys/tpw125 |
| LePT1 | *Lycopersicum esculentum* | CAA74607.1 | Daram et al. 1998 | https://doi.org/10.1007/s004250050394 |
| LePT2 | *Lycopersicum esculentum* | NP_001234043.1 | Liu et al. 1998 | https://doi.org/10.1104/pp.116.1.91 |
| LePT3 | *Lycopersicum esculentum* | NP_001318089.1 | Aoki et al. 2010 | https://doi.org/10.1186/1471-2164-11-210 |
| LePT4 | *Lycopersicum esculentum* | NP_001234674.2 | Nagy et al. 2005 | https://doi.org/10.1111/j.1365-313X.2005.02364.x |
| LePT6 | *Lycopersicum esculentum* | AJF19139.1 | Chen et al. 2014 | https://doi.org/10.1186/1471-2229-14-61 |
| LePT7 | *Lycopersicum esculentum* | AJF19140.1 | Chen et al. 2014 | https://doi.org/10.1186/1471-2229-14-61 |
| LePT8 | *Lycopersicum esculentum* | AJF19141.1 | Chen et al. 2014 | https://doi.org/10.1186/1471-2229-14-61 |
| LjPT3 | *Lotus japonicus* | BAE93353.1 | Maeda et al. 2006 | https://doi.org/10.1093/pcp/pcj069 |
| LjPT4 | *Lotus japonicus* | BAG71408.1 | Takeda et al. 2009 | https://doi.org/10.1111/j.1365-313X.2009.03824.x |
| MtPT1 | *Medicago truncatula* | AAB81346.1 | Liu et al. 1998 | https://doi.org/10.1094/MPMI.1998.11.1.14 |
| MtPT2 | *Medicago truncatula* | AAB81347.1 | Liu et al. 1998 | https://doi.org/10.1094/MPMI.1998.11.1.14 |
| MtPT3 | *Medicago truncatula* | ABM69110.1 | Liu et al. 1998 | https://doi.org/10.1094/MPMI.1998.11.1.14 |
| MtPT4 | *Medicago truncatula* | Q8GSG4.1 | Harrison et al. 2002 | https://doi.org/10.1105/tpc.004861 |
| MtPT5 | *Medicago truncatula* | ABM69111.1 | Liu et al. 1998 | https://doi.org/10.1094/MPMI.1998.11.1.14 |
| MtPT6 | *Medicago truncatula* | XP_003601529.1 | Breuillin-Sessoms et al. 2015 | https://doi.org/10.1105/tpc.114.131144 |
| MtPT7 | *Medicago truncatula* | A0A072UDN5 | Breuillin-Sessoms et al. 2015 | https://doi.org/10.1105/tpc.114.131144 |
| MtPT8 | *Medicago truncatula* | A0A072U3J3 | Breuillin-Sessoms et al. 2015 | https://doi.org/10.1105/tpc.114.131144 |
| MtPT9 | *Medicago truncatula* | A0A072VWQ9 | Breuillin-Sessoms et al. 2015 | https://doi.org/10.1105/tpc.114.131144 |
| MtPT10 | *Medicago truncatula* | A0A072VKI6 | Breuillin-Sessoms et al. 2015 | https://doi.org/10.1105/tpc.114.131144 |
| MtPT11 | *Medicago truncatula* | A0A072VWQ9 | Breuillin-Sessoms et al. 2015 | https://doi.org/10.1105/tpc.114.131144 |
| MtPT12 | *Medicago truncatula* | A0A072UDN5 | Breuillin-Sessoms et al. 2015 | https://doi.org/10.1105/tpc.114.131144 |
| MtPT13 | *Medicago truncatula* | A0A072U3J3 | Breuillin-Sessoms et al. 2015 | https://doi.org/10.1105/tpc.114.131144 |
| OsPT1 | *Oryza sativa* | Q8H6H4.1 | Paszkowski et al. 2002 | https://doi.org/0.1073/pnas.202474599 |
| OsPT2 | *Oryza sativa* | Q8GSD9.1 | Paszkowski et al. 2002 | https://doi.org/0.1073/pnas.202474599 |
| OsPT3 | *Oryza sativa* | Q7XDZ7.1 | Paszkowski et al. 2002 | https://doi.org/0.1073/pnas.202474599 |
| OsPT4 | *Oryza sativa* | Q8H6H2.1 | Paszkowski et al. 2002 | https://doi.org/0.1073/pnas.202474599 |
| OsPT5 | *Oryza sativa* | Q7X7V2.2 | Paszkowski et al. 2002 | https://doi.org/0.1073/pnas.202474599 |

| | | | | |
|---|---|---|---|---|
| OsPT6 | *Oryza sativa* | Q8H6H0.1 | Paszkowski et al. 2002 | https://doi.org/0.1073/pnas.202474599 |
| OsPT7 | *Oryza sativa* | Q8H6G9.1 | Paszkowski et al. 2002 | https://doi.org/0.1073/pnas.202474599 |
| OsPT8 | *Oryza sativa* | Q8H6G8.1 | Paszkowski et al. 2002 | https://doi.org/0.1073/pnas.202474599 |
| OsPT9 | *Oryza sativa* | Q8H6G7.2 | Paszkowski et al. 2002 | https://doi.org/0.1073/pnas.202474599 |
| OsPT10 | *Oryza sativa* | Q69T94.1 | Paszkowski et al. 2002 | https://doi.org/0.1073/pnas.202474599 |
| OsPT11 | *Oryza sativa* | Q94DB8.1 | Paszkowski et al. 2002 | https://doi.org/0.1073/pnas.202474599 |
| PtPT10 | *Populus trichocarpa* | XP_006374329 | Loth-Pereda et al. 2011 | https://doi.org/10.1104/pp.111.180646 |
| SbPT1 | *Sorghum bicolor* | NP_001275200.1 | Leggewie et al. 1997 | https://doi.org/10.1105/tpc.9.3.381 |
| SbPT2 | *Sorghum bicolor* | CAA67396.1 | Leggewie et al. 1997 | https://doi.org/10.1105/tpc.9.3.381 |
| SbPT3 | *Sorghum bicolor* | CAC87043.1 | Rausch et al. 2001 | https://doi.org/10.1038/35106601 |
| SbPT4 | *Sorghum bicolor* | AAW51149.1 | Karandashov and Bucher, 2005 | https://doi.org/10.1016/j.tplants.2004.12.003 |
| SbPT5 | *Sorghum bicolor* | AAX85195.1 | Nagy et al. 2005 | https://doi.org/10.1111/j.1365-313X.2005.02364.x |
| SbPT11 | *Sorghum bicolor* | XP_002458253.1 | Walder et al. 2015 | https://doi.org/10.1111/nph.13292 |
| TaPT1 | *Triticum aestivum* | CAC69856.1 | Davies et al. 2002 | https://doi.org/10.1046/j.1365-3040.2002.00913.x |
| TaPT8 | *Triticum aestivum* | AAP49822.1 | Chang et al. 2003 | unpublished |
| TmPT1 | *Triticum monococcum* | AAQ06280.1 | Ma et al. 2002 | unpublished |
| VvPT1 | *Vitis vinifera* | XP_002267369.1 | Valat et al. 2018 | https://doi.org/10.1007/s00572-017-0809-5 |
| VvPT2 | *Vitis vinifera* | XP_002267327.1 | Valat et al. 2018 | https://doi.org/10.1007/s00572-017-0809-5 |
| ZmPT1 | *Zea mays* | NP_001105269.2 | Wright et al. 2005 | https://doi.org/10.1111/j.1469-8137.2005.01472.x |
| ZmPT2 | *Zea mays* | NP_001105816.1 | Schnable et al. 2009 | https://doi.org/10.3390/ijms17060930 |
| ZmPT3 | *Zea mays* | AAY42387.1 | Nagy et al. 2005 | unpublished |
| ZmPT4 | *Zea mays* | AAY42388.1 | Nagy et al. 2005 | unpublished |
| ZmPT5 | *Zea mays* | ACG37120.1 | Alexendrov et al. 2009 | https://doi.org/10.1007/s11103-008-9415-4 |
| ZmPT6 | *Zea mays* | NP_001105776.1 | Liu et al. 2016 | https://doi.org/10.3390/ijms17060930 |
